# R-pyocin-mediated selection restores aminoglycoside sensitivity in a pan-drug resistant *Pseudomonas aeruginosa* isolate

**DOI:** 10.64898/2026.08.04.742823

**Authors:** Isaac Estrada, Davina Campbell, Garrett S. Welch, Parker Smith, Joanna B. Goldberg, Kendra P. Rumbaugh, Stephen P. Diggle

## Abstract

The global escalation of antibiotic resistance is a critical threat necessitating innovative strategies to provide new therapeutic options and restore the efficacy of conventional drugs. *Pseudomonas aeruginosa* exemplifies this challenge by utilizing a robust genomic resistome to persist in clinical settings. Here, we demonstrate that R-pyocins (phage-like bactericidal particles) can be leveraged not as conventional biocides but as precise selective forces to drive an evolutionary trap. We subjected the laboratory strains PAO1 and PAK and the clinical pan-drug-resistant (PDR) wound isolate MRSN 6220 to R-pyocin selective pressure. Under sustained R-pyocin pressure, every pyomelanogenic variant we recovered evaded R-pyocin killing through large-scale chromosomal deletions spanning 250-388 kbp. These deletions encompass a conserved region harboring the *galU* gene (essential for LPS synthesis), the *hmgA* gene (yielding a pyomelanogenic ‘brown’ phenotype), and the *mexXYZ* multidrug efflux operon. While the loss of *galU* confers broad cross-resistance to R-pyocins by likely truncating the LPS receptor, the concurrent excision of *mexXY* induces profound collateral sensitivity to aminoglycosides. These deletions also eliminate critical virulence factors and biofilm clusters, including the *hcnABC*, *exoY*, *phzABCDEFG*, and *cup* operons. In a *Galleria mellonella* model, the resulting brown mutants were non-lethal, and in infected murine wound tissue treated *ex vivo* with gentamicin they exhibited a significant 3-log reduction in bacterial load, whereas the pan-drug-resistant ancestor did not. This work establishes a framework for utilizing R-pyocins as evolutionary steering agents that couple the acquisition of R-pyocin resistance to the reversion of multidrug resistance into an attenuated, biofilm-deficient, and clinically manageable state.

**Importance:** Pan-drug-resistant (PDR) pathogens demand novel strategies that both kill and restore antibiotic efficacy. Here, we describe an evolutionary trap for *Pseudomonas aeruginosa*, where selection for R-pyocin resistance drives large-scale (∼300 kb) chromosomal remodeling. Although these deletions confer R-pyocin immunity via loss of the *galU* gene, they simultaneously collapse the pathogen’s virulence and defense. Crucially, the excision of the *mexXY* efflux operon resensitizes a PDR clinical isolate to conventional aminoglycosides, while the collateral loss of critical virulence and biofilm clusters abrogates pathogenesis. By coupling resistance acquisition to substantial fitness costs, our work establishes a framework for using R-pyocins to force predictable evolutionary trade-offs, driving the reversion of multidrug resistance to an attenuated, biofilm-deficient, and clinically manageable state.

## Introduction

The global rise of antibiotic resistance represents one of the most critical threats to modern medicine, rendering standard treatments ineffective and resulting in once-manageable infections turning deadly (1). As conventional antibiotics fail, the evolutionary adaptability of bacterial pathogens has outpaced drug development, creating an urgent need for alternative therapeutic paradigms (2). The global dissemination of multi-(MDR), extensively-(XDR), and pan-drug (PDR) resistant *Pseudomonas aeruginosa* represents a critical inflection point in clinical microbiology, where traditional small-molecule therapeutics increasingly fail to provide clinical resolution (3, 4). *P. aeruginosa* leverages a sophisticated, multifaceted resistome, primarily driven by the synergistic action of low outer membrane permeability and the expression of broad-spectrum multidrug efflux pumps, such as the Resistance-Nodulation-Division (RND) family (3, 5, 6). Among these, the MexXY-OprM system is a principal determinant of resistance to aminoglycosides and is frequently upregulated via mutations in the transcriptional repressor MexZ (7, 8). As the pipeline for novel antibiotics remains constrained, there is an emerging need for alternative antimicrobial modalities that can either kill resistant bacteria and/or systematically circumvent established resistance mechanisms.

R-pyocins are high-molecular-weight contractile bacteriocins produced by *P. aeruginosa* that have emerged as potent candidates for targeted antimicrobial therapy (9–11). These evolutionarily conserved structures are co-opted phage tail-like nano-machines, which are genetically and morphologically analogous to the tails of P2-like *Myoviridae* bacteriophage (12–14). R-pyocins belong to the broader class of contractile tailocins, phage-derived bactericidal nanomachines produced across diverse bacterial genera, and tailocin is increasingly the preferred term for these particles. We retain R-pyocin throughout to denote the *P. aeruginosa*-specific subtypes examined here. The lethality of R-pyocins is characterized by a highly efficient, single-hit killing kinetic: upon specific recognition of the host’s lipopolysaccharide (LPS) core, the sheath undergoes a rapid conformational contraction, driving a rigid inner tube through the cell envelope to depolarize the cytoplasmic membrane (15–17). While a substantial body of work has elucidated the structural and biophysical requirements for R-pyocin-mediated killing (18–20), the specific genetic pathways through which *P. aeruginosa* evolves resistance to these bacteriocins, and the subsequent evolutionary trade-offs imposed by such resistance, remain poorly understood.

Previous studies have established that *P. aeruginosa* resists phage predation through highly plastic defense strategies, including CRISPR-Cas systems (21–23) and large chromosomal deletions (LCDs) mediated by MutL and non-homologous end joining (24, 25). While these mechanisms demonstrate the capacity for bacterial genome remodeling in response to selection, they often result in a heterogeneous, stochastic population and diverse resistance profiles that are difficult to manage clinically. Further, unlike lytic phages, which exert multifaceted and often unpredictable selective pressures (22, 23), R-pyocins provide a targeted, single-hit kinetic challenge (13, 26). We hypothesized that this precision could drive a more predictable evolutionary trajectory toward antibiotic sensitization and virulence attenuation. Characterizing whether R-pyocin-mediated selection forces a predictable genomic reorganization is essential for determining if these bacteriocins offer a reliable means of forcing the reversion of multidrug resistance in clinical settings.

In this study, we investigated the evolutionary trajectories of laboratory and clinical PDR *P. aeruginosa* under R-pyocin-mediated selective pressure. We demonstrate that R-pyocin resistant variants recovered under this selection regime consistently harbor extensive (250–388 kbp) chromosomal deletions, an evolutionary trap that couple’s survival to two linked trade-offs. The excision of the *galU* gene results in R-pyocin cross-resistance and is inextricably linked to the loss of the *mexXY* multidrug efflux operon and multiple critical virulence and biofilm-forming loci (including *hcnABC*, *exoY*, *phzABCDEFG*, and *cup* operons). This genomic remodeling renders a PDR strain sensitive to aminoglycosides in a process we term pyocin-mediated collateral sensitivity (PMCS), abolishes biofilm formation, attenuates virulence in a *Galleria mellonella* model and significantly enhances the effectiveness of antibiotics in a murine infection model. By coupling the survival of a bacteriocin challenge to the systematic dismantling of the pathogen’s defensive and offensive machinery, we establish a framework for using R-pyocins as potent evolutionary steering agents, forcing the reversion of a recalcitrant, pan-drug resistant strain into an attenuated, manageable state.

## Results

### Evolutionary selection of R-pyocin resistance induces a conserved pyomelanogenic phenotype and broad R-pyocin cross-resistance

To investigate the evolutionary trajectories associated with R-pyocin resistance, we subjected the laboratory strains PAO1 and PAK, and a pan-drug resistant (PDR) clinical isolate MRSN 6220 each to selective pressure from varying concentrations (0.098 to 100 µg/mL) of 2 R-pyocin subtypes to which they exhibited susceptibility (**Fig. 1A**). PAO1 was exposed to an R1-pyocin produced by strain MRSN 2101 and an R5-pyocin produced by MRSN 1344. PAK was exposed to an R2-pyocin produced by strain PAO1 and an R5-pyocin produced by MRSN 1344. MRSN 6220 was likewise exposed to an R2-pyocin produced by PAO1 and the R5-pyocin produced by MRSN 1344 (**Table 1**). Negative control R-pyocins were also used, in which the receptor binding domain of the R-pyocin tail fiber had been deleted, as previously described (27, 28). Exposure to any active R-pyocin subtype inhibited normal growth of the bacterial cultures in the first 24 hrs. At 48 h post treatment, the emergence of R-pyocin resistant isolates characterized by a distinct hypermelanogenic “brown” (Brn) pigmented phenotype became apparent (**Fig. 1B, C**). Initially we focused on two Brn mutants from both PAO1 and 6220 (PAO1-1A2, PAO1-1C1, 6220-2A3, and 6220-2G2) and retested for R-pyocin susceptibility. All Brn mutants showed complete cross-resistance to all R-pyocins to which its ancestor strain was previously susceptible (**Fig. 1D**).

**Figure 1:**
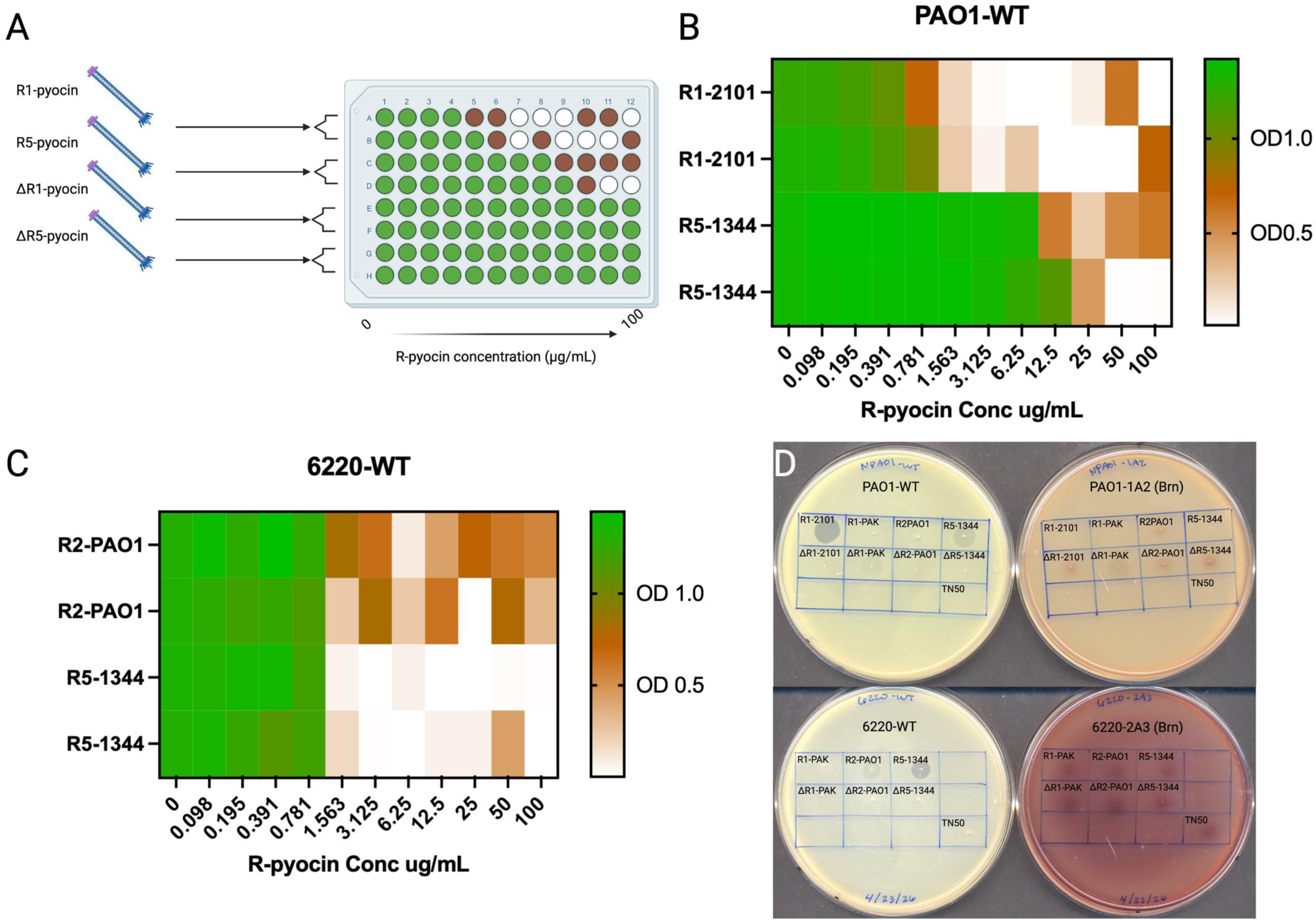
Selection and phenotypic characterization of R-pyocin resistant mutants. **(A)** Schematic overview of the broth microdilution selection approach used to isolate resistant populations under a gradient of R-pyocin concentrations ranging from 0 to 100 µg/mL. The color-coded matrix illustrates the distribution of distinct breakthrough phenotypes emerging during selective challenge. Green wells indicate uninhibited bacterial growth, brown wells indicated breakthrough-resistant phenotypes, and white wells indicate total bacterial inhibition. **(B)** Heatmap of the R-pyocin inhibition profile of PAO1-WT across a range of concentrations for R1-2101 and R5-1344 pyocins. Green regions indicate wells where robust bacterial density was observed, brown regions indicate well where inhibited bacterial density and the brown phenotype was observed, and white regions indicated wells where total bacterial inhibition was observed. **(C)** Heatmap of the R-pyocin inhibition profile of 6220-WT across a range of concentrations for R2-PAO1 and R5-1344 pyocins. Color-coding is the same as previously stated. **(D)** Spot assays of ancestor strains PAO1-WT and 6220-WT and their derived R-pyocin resistant Brn mutants, PAO1-1A2 and 6220-2A3. Each strain was challenged with each of the R-pyocin subtypes: R1-2101, R1-PAK, R2-PAO1 and R5-1344. Additionally, a R-pyocin knockout was generated for each of the R-pyocin subtypes, notated by Δ. These strains have had the receptor binding region of the R-pyocin deleted, attenuating the tailocin’s ability to bind to LPS receptors. Spot assays of the Brn mutants show that this phenotype develops cross-resistance to all R-pyocin subtypes, regardless of the ancestral strain susceptibility.

**Table 1:**
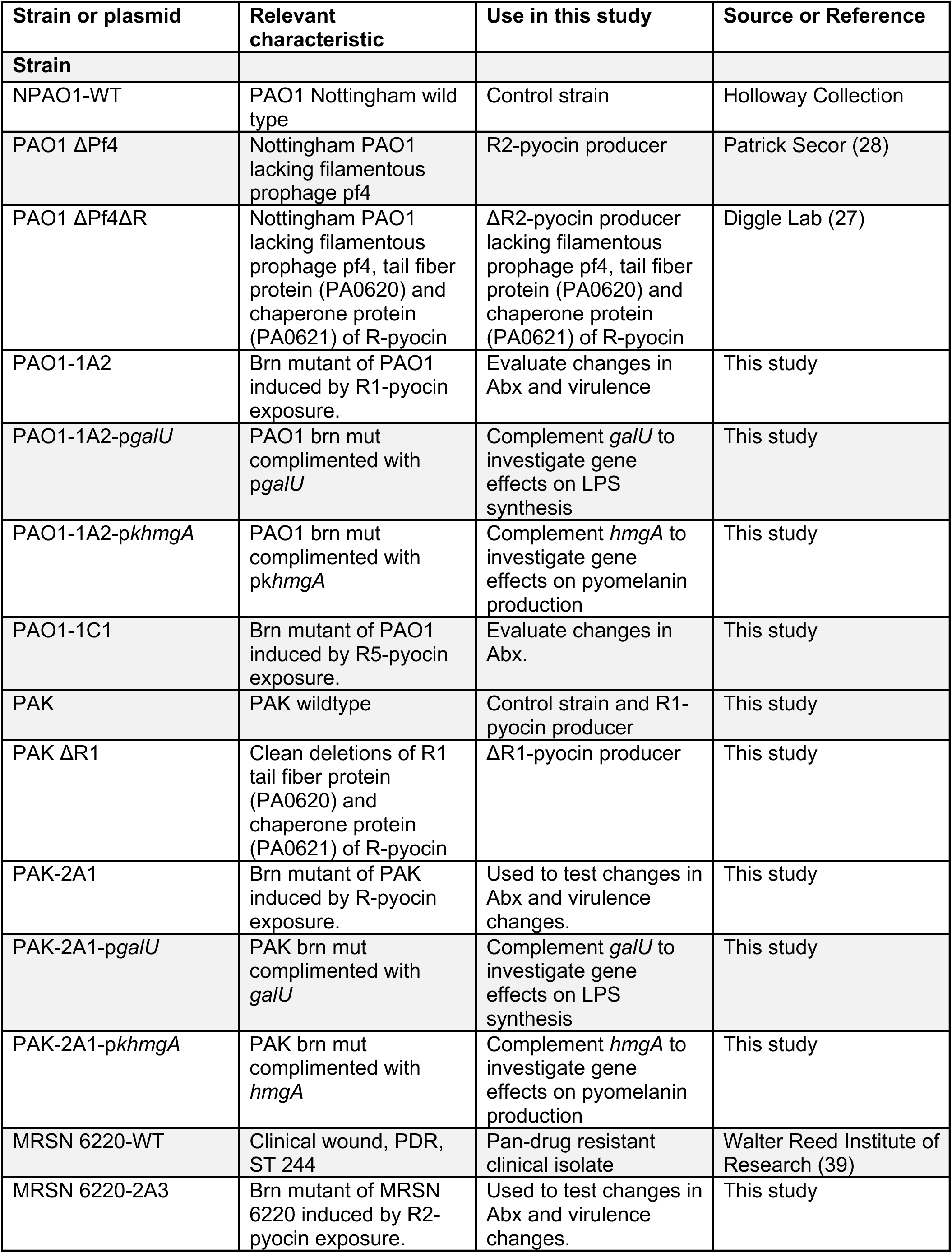
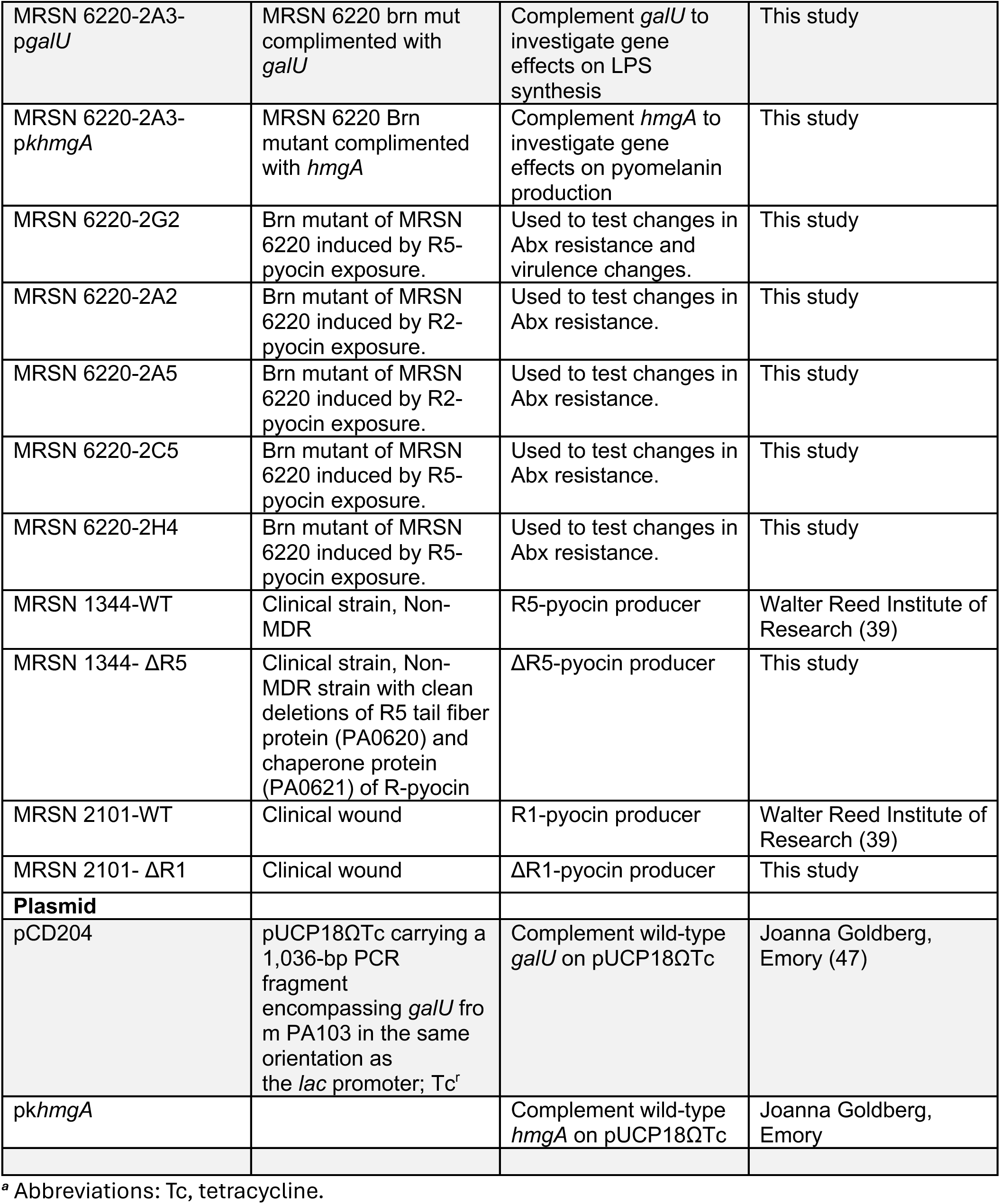
Lists the bacterial strains and plasmids used in this study and their sources.

| Strain or plasmid | Relevant characteristic | Use in this study | Source or Reference |
|---|---|---|---|
| **Strain** |  |  |  |
| NPAO1-WT | PAO1 Nottingham wild type | Control strain | Holloway Collection |
| PAO1 ΔPf4 | Nottingham PAO1 lacking filamentous prophage pf4 | R2-pyocin producer | Patrick Secor (28) |
| PAO1 ΔPf4ΔR | Nottingham PAO1 lacking filamentous prophage pf4, tail fiber protein (PA0620) and chaperone protein (PA0621) of R-pyocin | ΔR2-pyocin producer lacking filamentous prophage pf4, tail fiber protein (PA0620) and chaperone protein (PA0621) of R-pyocin | Diggle Lab (27) |
| PAO1-1A2 | Brn mutant of PAO1 induced by R1-pyocin exposure. | Evaluate changes in Abx and virulence | This study |
| PAO1-1A2-p*galU* | PAO1 brn mut complimented with p*galU* | Complement *galU* to investigate gene effects on LPS synthesis | This study |
| PAO1-1A2-p*khmgA* | PAO1 brn mut complimented with pk*hmgA* | Complement *hmgA* to investigate gene effects on pyomelanin production | This study |
| PAO1-1C1 | Brn mutant of PAO1 induced by R5-pyocin exposure. | Evaluate changes in Abx. | This study |
| PAK | PAK wildtype | Control strain and R1-pyocin producer | This study |
| PAK ΔR1 | Clean deletions of R1 tail fiber protein (PA0620) and chaperone protein (PA0621) of R-pyocin | ΔR1-pyocin producer | This study |
| PAK-2A1 | Brn mutant of PAK induced by R-pyocin exposure. | Used to test changes in Abx and virulence changes. | This study |
| PAK-2A1-p*galU* | PAK brn mut complimented with *galU* | Complement *galU* to investigate gene effects on LPS synthesis | This study |
| PAK-2A1-p*khmgA* | PAK brn mut complimented with *hmgA* | Complement *hmgA* to investigate gene effects on pyomelanin production | This study |
| MRSN 6220-WT | Clinical wound, PDR, ST 244 | Pan-drug resistant clinical isolate | Walter Reed Institute of Research (39) |
| MRSN 6220-2A3 | Brn mutant of MRSN 6220 induced by R2-pyocin exposure. | Used to test changes in Abx and virulence changes. | This study |
| MRSN 6220-2A3-p*galU* | MRSN 6220 brn mut complimented with *galU* | Complement *galU* to investigate gene effects on LPS synthesis | This study |
| MRSN 6220-2A3-p*khmgA* | MRSN 6220 Brn mutant complimented with *hmgA* | Complement *hmgA* to investigate gene effects on pyomelanin production | This study |
| MRSN 6220-2G2 | Brn mutant of MRSN 6220 induced by R5-pyocin exposure. | Used to test changes in Abx resistance and virulence changes. | This study |
| MRSN 6220-2A2 | Brn mutant of MRSN 6220 induced by R2-pyocin exposure. | Used to test changes in Abx resistance. | This study |
| MRSN 6220-2A5 | Brn mutant of MRSN 6220 induced by R2-pyocin exposure. | Used to test changes in Abx resistance. | This study |
| MRSN 6220-2C5 | Brn mutant of MRSN 6220 induced by R5-pyocin exposure. | Used to test changes in Abx resistance. | This study |
| MRSN 6220-2H4 | Brn mutant of MRSN 6220 induced by R5-pyocin exposure. | Used to test changes in Abx resistance. | This study |
| MRSN 1344-WT | Clinical strain, Non-MDR | R5-pyocin producer | Walter Reed Institute of Research (39) |
| MRSN 1344- ΔR5 | Clinical strain, Non-MDR strain with clean deletions of R5 tail fiber protein (PA0620) and chaperone protein (PA0621) of R-pyocin | ΔR5-pyocin producer | This study |
| MRSN 2101-WT | Clinical wound | R1-pyocin producer | Walter Reed Institute of Research (39) |
| MRSN 2101- ΔR1 | Clinical wound | ΔR1-pyocin producer | This study |
| **Plasmid** |  |  |  |
| pCD204 | pUCP18ΩTc carrying a 1,036-bp PCR fragment encompassing *galU* from PA103 in the same orientation as the *lac* promoter; Tc<sup>r</sup> | Complement wild-type *galU* on pUCP18ΩTc | Joanna Goldberg, Emory (47) |
| pk*hmgA* |  | Complement wild-type *hmgA* on pUCP18ΩTc | Joanna Goldberg, Emory |

<sup>*a*</sup> Abbreviations: Tc, tetracycline.

### Genomic analysis of R-pyocin-resistant mutants reveals a conserved 250-kbp deletion region

To resolve the genetic basis of R-pyocin resistance and its associated collateral effects, we performed hybrid assembly-based genomic analysis on the five Brn mutants spanning three strain backgrounds (**Fig. 2A, B; Tables S1 to S5**). We identified a conserved, large-scale chromosomal deletion of approximately 250 kb in all Brn variants, regardless of strain background. This core deletion region encompassed 175 genes shared among all five sequenced mutants, including the *mexXY* operon and its transcriptional regulator *mexZ*. The deletion also included the *galU* and *hmgA* genes. Beyond resistance and metabolic markers, this 175-gene loss resulted in the elimination of several well-established virulence factors, including the hydrogen cyanide synthase cluster (*hcnABC*), the Type III secretion effector *exoY*, and the surface structure and biofilm-associated *cupA* fimbrial subunits. While the core 250-kb deletion was conserved, we observed notable differences in the total extent of chromosomal loss between the three strain backgrounds. Mutants derived from PAO1 (PAO1-1A2 and PAO1-1C1) exhibited an expanded deletion totaling 388.4 kb, followed by the PAK-derived mutant PAK-2A1 at 364.4 kb (**Table S5**). The regions extending beyond the conserved core in these two backgrounds included loci contributing to additional surface structure and biofilm formation, specifically the functional amyloid (*fap*) operon, the pyrroloquinoline quinone (*pqqABCDEF*) biosynthetic cluster, and the biofilm dispersal regulator *bdlA*, none of which were lost in the MRSN 6220-derived mutants. The phenazine biosynthetic cluster (*phzABCDEFG*) was deleted only in the PAO1 lineage. Additional deletions in the MRSN 6220-derived mutants (6220-2A3 and 6220-2G2) were exclusively related to prophage loci, indicating prophage excision activity under R-pyocin-mediated selective pressure. Notably, the conserved region also encompassed *ku* and *ligD* in all five mutants, which encode the non-homologous end joining machinery implicated in the generation of large chromosomal deletions in *P. aeruginosa* (24, 25). A list of common deleted genes and functions across all Brn mutants is available in **Table 2**. A full list of all gene deletions across the isolated variants is available in **Tables S1 to S5**, organized sequentially by strain background (Table S1: PAO1-1A2; Table S2: PAO1-1C1; Table S3: 6220-2A3; Table S4: 6220-2G2; Table S5: PAK-2A1).

**Figure 2:**
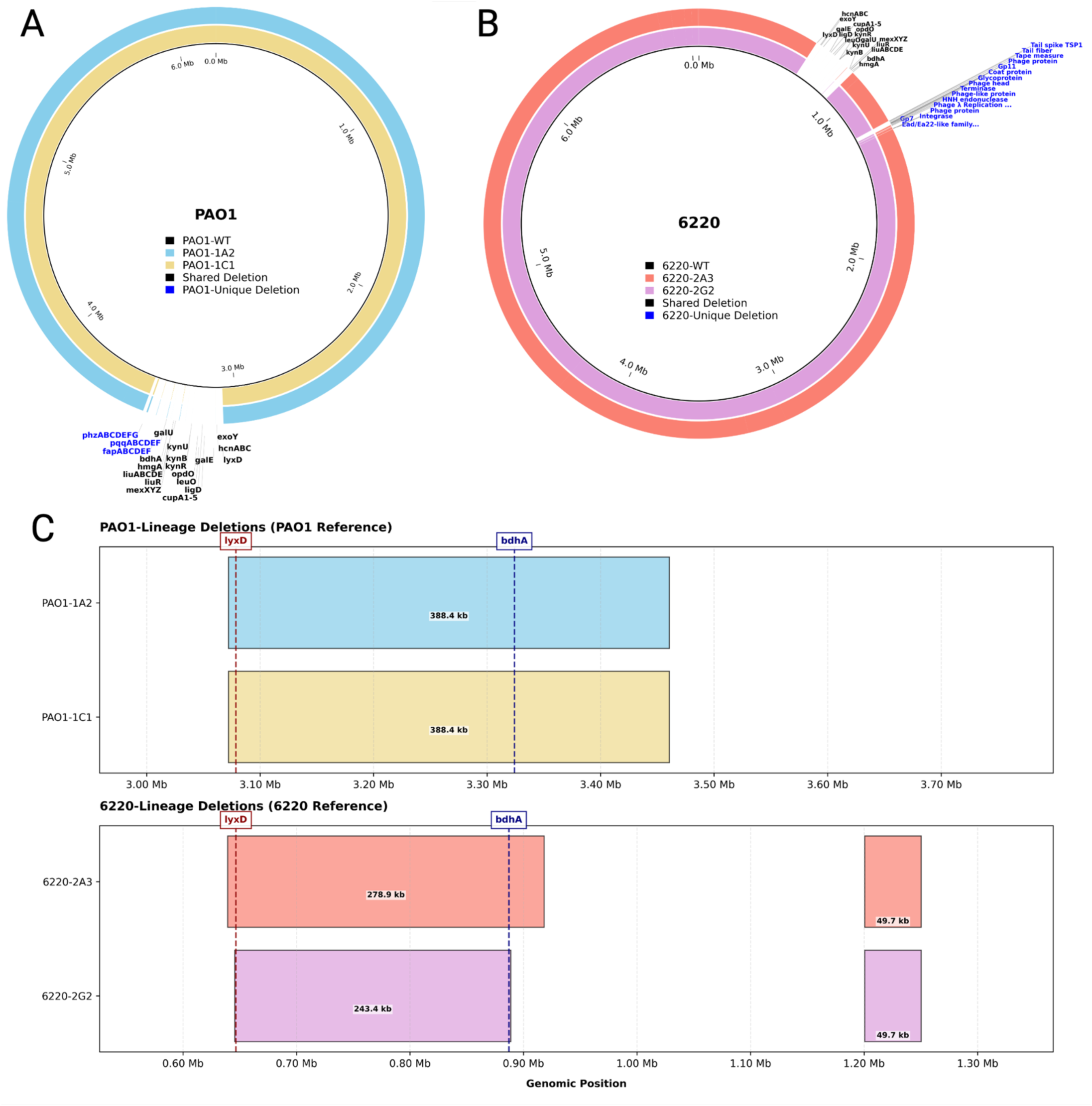
R-pyocin resistance is mediated by large chromosomal deletions. **(A)** Circular genome map of PAO1 Brn mutants, PAO1-1A2 and PAO1-1C1 compared to the PAO1-WT ancestor. Chromosomal deletion regions are identical in both R-pyocin resistant Brn mutants and show loss of specific genetic clusters, including the *mexXYZ* operon, *galU*, and the hydrogen cyanide operon *hcnABC*, as well as the phenazine biosynthesis operon and fimbrial subunits *cupA1-5*. **(B)** Circular genome map of PDR MRSN 6220 and R-pyocin resistant Brn mutants, 6220-2A3 and 6220-2G2. Labels highlight gene deletions found in the Brn mutants, such as *mexXYZ* operon, *galU*, and the hydrogen cyanide operon *hcnABC*. Multiple prophage-related genes are also noted. Black gene labels identify gene deletions found in all 4 Brn mutants. Blue gene labels identify gene deletions unique to the strain background, PAO1 and 6220. **(C)** A linear map of the gene deletions found in 4 Brn mutants. Both PAO1 and 6220-derived mutants exhibit chromosomal reorganizations, with a conserved region spanning approximately from *lyxD* to the *bdhA* gene. While PAO1-lineage mutants show a single continuous 388.4kb deletion, 6220-lineage mutants display a similar deletion region, as well as discrete, prophage-related deletion regions. The PAK-derived mutant PAK-2A1 was also sequenced but is not shown here; its deletion spans 364.4 kb and is reported in Table S5.

**Table 2.** List of common gene deletions found in all five Brn mutants due to Pyocin-Mediated Collateral Sensitivity (PMCS).

| **Gene name** | **Product** | **Function** |
|---|---|---|
| *lyxD* | L-lyxonate dehydratase | Carbohydrate Metabolism |
| *hcnABC* | cyanide-forming glycine dehydrogenase subunits HcnA, HcnB and HcnC | Virulence Factor |
| *exoY* | type III secretion system effector, ExoY adenylate cyclase | Virulence Factor |
| *clpP* | Clp protease | Protein Quality Control |
| *katE* | catalase HPII | Oxidative Stress Defense |
| *gor* | glutathione-disulfide reductase | Oxidative Stress Defense |
| *galE* | NAD-dependent epimerase | Carbohydrate Metabolism |
| *ligD* | DNA ligase D | DNA Repair |
| *cupA1-5* | Components of the cupA chaperone-usher fimbrial assembly system | Adhesion & Biofilm Formation |
| *leuO* | HTH-type transcriptional regulator LeuO | Transcriptional Regulation |
| *opdO* | Porin | Membrane Transport |
| *kynR* | kynurenine pathway transcriptional regulator KynR | Transcriptional Regulation |
| *kynB* | kynurenine formamidase KynB | Amino Acid Catabolism |
| *kynU* | kynureninase | Amino Acid Catabolism |
| *copA* | Copper resistance protein A | Heavy Metal Resistance |
| *copB* | Copper resistance protein B | Heavy Metal Resistance |
| *cynR* | transcriptional regulator CynR | Transcriptional Regulation |
| *cynT* | carbonic anhydrase CynT | Cyanate Metabolism / Detoxification |
| *cynS* | cyanase | Detoxification |
| *galU* | UTP--glucose-1-phosphate uridylyltransferase GalU | Cell Envelope Biosynthesis |
| *mexZ* | TetR family transcriptional regulator AmrR | Antibiotic Resistance Regulation |
| *mexX* | multidrug efflux RND transporter periplasmic adaptor subunit MexX | Multidrug Efflux |
| *mexY* | multidrug efflux RND transporter permease subunit MexY | Multidrug Efflux |
| *liuR* | liu genes transcriptional regulator LiuR | Transcriptional Regulation |
| *liuABCDE* | liu cluster involved in leucine and acyclic terpene utilization | Amino Acid / Terpene Catabolism |
| *hmgA* | homogentisate 1,2-dioxygenase | Amino Acid Catabolism |
| *bdhA* | 3-hydroxybutyrate dehydrogenase | Ketone Body Metabolism |

### Large-scale chromosomal deletions drive R-pyocin resistance, biofilm impairment, and virulence attenuation

To confirm that disruption of the LPS core biosynthesis pathway drives R-pyocin resistance, while the brown phenotype results from the concurrent deletion of metabolic loci within the excised chromosomal region, we performed targeted complementation assays on the brown mutants isolated from the PAO1, PAK, and MRSN 6220 lineages. We found that introducing a wild-type copy of *galU* on a plasmid successfully reverted the isolates MRSN 6220-Brn and PAK-Brn, restoring their susceptibility to wild-type R-pyocins (**Fig. 3A, B**). Intriguingly, however, *galU* expression failed to restore R-pyocin susceptibility in our PAO1-Brn strain. This striking divergence indicates that while *galU* disruption is a route to R-pyocin resistance, our PAO1 background relies on additional or alternative genetic factors within its specific large-scale chromosomal deletion boundary (or other genetic factors) to mediate functional R-pyocin targeting. As expected, complementation with *hmgA* restored wild-type pigmentation to all Brn mutants, confirming that the pyomelanogenic phenotype resulted from *hmgA* disruption (**Fig. 3A**).

**Figure 3:**
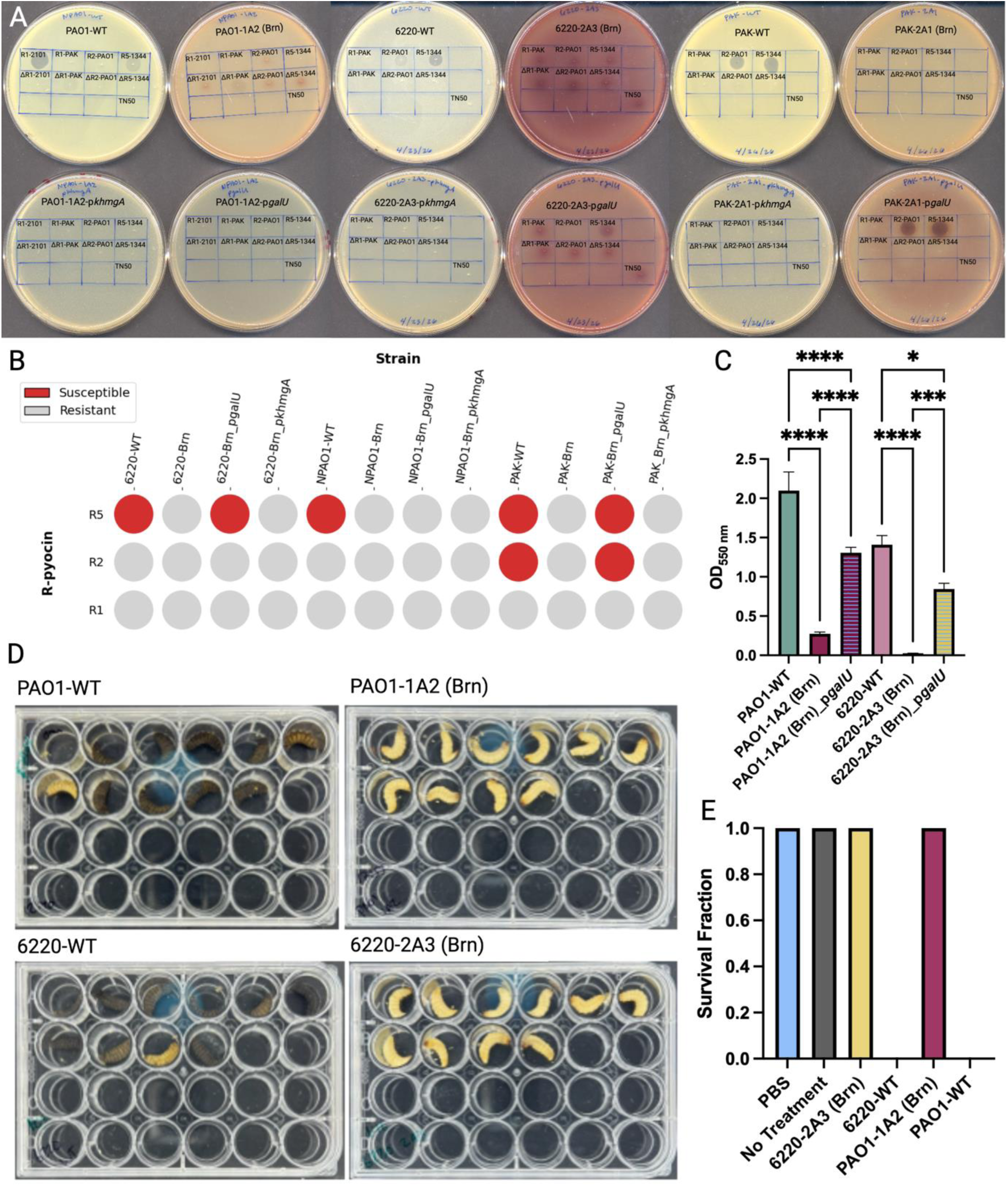
Deletion of *galU* drives R-pyocin resistance and attenuated biofilm formation and virulence. **(A)** Spot assay of PAO1-WT, 6220-WT, and PAK-WT, Brn mutant derivatives, and complemented versions carrying *hmgA* or *galU*. Brn phenotype mutants are cross-resistant to all R-pyocin subtypes, while complementation with *galU* restores R-pyocin sensitivity to the 6220 and PAK Brn mutants, but not the PAO1 Brn mutant. Complementation with *hmgA* reverses the pyomelanin overproduction. **(B)** Dotplot summary and R-pyocin activity against PAO1, 6220 and PAK lineage strains. Red circles indicate susceptibility and grey circles indicate resistance. **(C)** Biofilm quantification of ancestor strains, Brn mutants, and *galU* complements determined via crystal violet staining. Complementation with *galU* partially restores ability of Brn mutants to attach and form biofilms. Sidak’s multiple comparison test. (*) p = 0.0436. (***) p = 0.0009, (****) p < 0.0001. Error bars = SEM. **(D)** Visual assessment of *Galleria mellonella* larvae 24-hours post-injection with the ancestor strains PAO1-WT and 6220-WT, undergo widespread melanization and mortality, while those injected with Brn mutants, PAO1-1A2 and 6220-2A3, remain healthy. **(E)** Survival fraction of *G. mellonella* larvae monitored over a 24-hour period. Larvae infected with the Brn mutants, 6220-2A3 and PAO1-1A2, exhibit 100% survival, identical to the PBS and No-Treatment control groups. Ancestor strains PAO1-WT and 6220-WT result in complete larvae mortality within the same incubation period.

Because the large-scale chromosomal deletions in the Brn mutants excised multiple established biofilm-related loci (including *galU*, the *cupA1-5* fimbrial subunits, and *bdlA*), we reasoned that biofilm architecture would be severely compromised. To test this hypothesis, we evaluated the biofilm-forming capacity of the wild-type ancestors (PAO1-WT and 6220-WT) and their respective Brn mutants (PAO1-1A2 and 6220-2A3) using crystal violet biofilm assays. As anticipated, both Brn mutants exhibited a profound and significant reduction in biofilm formation compared to their wild-type ancestors (**Fig. 3C**). Because *galU* encodes a key enzyme producing UDP-glucose (a vital precursor for the synthesis of matrix exopolysaccharides), we sought to determine if introducing a wild-type copy of *galU* in trans could rescue this phenotype. Complementation with *galU* resulted in a significant, yet incomplete, restoration of biofilm attachment in both the PAO1-1A2 and 6220-2A3 backgrounds (**Fig. 3C**). While restoring *galU* replenished critical biosynthetic precursors, it did not fully compensate for the multi-locus deletion of the other core structural and regulatory biofilm genes lost in the 250 to 388 kbp chromosomal excisions.

Beyond compromising biofilm architecture, the chromosomal deletions identified in the Brn mutants strongly pointed toward attenuation of pathogenic potential. Genomic mapping revealed that this excised region contained a suite of well-characterized, essential virulence factors (**Fig. 2)**. Most notably, all five Brn mutants lacked the hydrogen cyanide synthase (*hcnABC*) operon (a key mediator of acute cytotoxicity) and *exoY*, a potent Type III secretion system effector toxin. Given the simultaneous loss of these acute toxins alongside the critical surface-structure and biofilm-related machinery, we hypothesized that these mutants would be profoundly impacted in their ability to kill a host. To validate this hypothesized loss of pathogenicity *in vivo*, we evaluated the mutants using a *Galleria mellonella* (wax moth larvae) infection model (**Fig. 3D**). The wild-type ancestors, PAO1-WT and MRSN 6220-WT, demonstrated acute, aggressive lethality, causing complete melanization and larval mortality within 24 hours (**Fig. 3E**). Conversely, Brn mutants were rendered entirely non-lethal, yielding a 1.0 survival fraction that matched the sterile PBS and no-treatment negative controls (**Fig. 3E**). This abolition of *in vivo* virulence demonstrates evolutionary trade-offs and biological fitness costs imposed on *P. aeruginosa* when it becomes resistant to R-pyocins through large-scale genomic deletions, effectively converting a pan-drug resistant pathogen into an avirulent state.

### R-pyocin-mediated collateral sensitivity restores aminoglycoside sensitivity and gentamicin activity in a *P. aeruginosa* wound infection

We next evaluated whether the chromosomal deletions driving R-pyocin resistance incurred pleiotropic fitness costs that could be exploited therapeutically. Using Kirby-Bauer disk diffusion assays (29–31), to profile the mutants against a broad panel of antimicrobials, we identified striking instances of collateral sensitivity. Because the standard laboratory strain PAO1 is inherently antibiotic-sensitive, the pan-drug resistant (PDR) clinical isolate MRSN 6220 provided a highly relevant genetic background to evaluate the restoration of drug susceptibility. Notably, all six 6220-Brn mutants exhibited substantially increased zones of inhibition for the aminoglycosides amikacin (AK), gentamicin (CN) and tobramycin (TOB), while zones for the β-lactams and fluoroquinolones were unchanged (Fig. 4A, B, Table S6). The magnitude of these shifts was large, with the amikacin zone increasing from 7.3 mm for 6220-WT to between 22.0 and 23.8 mm across the mutants, and the gentamicin zone from 10.4 mm to between 23.8 and 27.2 mm. Current CLSI criteria do not assign interpretive breakpoints for gentamicin against *P. aeruginosa* and restrict amikacin interpretation to urinary isolates (31), so these shifts cannot be expressed as a change in susceptibility category. For tobramycin, which retains interpretive criteria, zones increased from no measurable inhibition for 6220-WT to between 11.0 and 15.5 mm, moving five of six mutants from the resistant into the intermediate category. We therefore report these data as a substantial and mechanistically explained reduction in aminoglycoside resistance rather than as a categorical reversion to clinical susceptibility, and we assess the functional consequence of this shift directly by gentamicin challenge of infected wound tissue (Fig. 5). This phenotypic reversal aligns with our genomic data, which revealed that the large-scale chromosomal excision included the *mexXYZ* operon, disabling the MexXY-OprM multidrug efflux pump, which is a primary determinant of aminoglycoside resistance in *P. aeruginosa*. We term this mechanism Pyocin-Mediated Collateral Sensitivity (PMCS). The term denotes the selection context examined here and is not intended to assign the underlying deletion-associated mechanism exclusively to R-pyocins.

**Figure 4:**
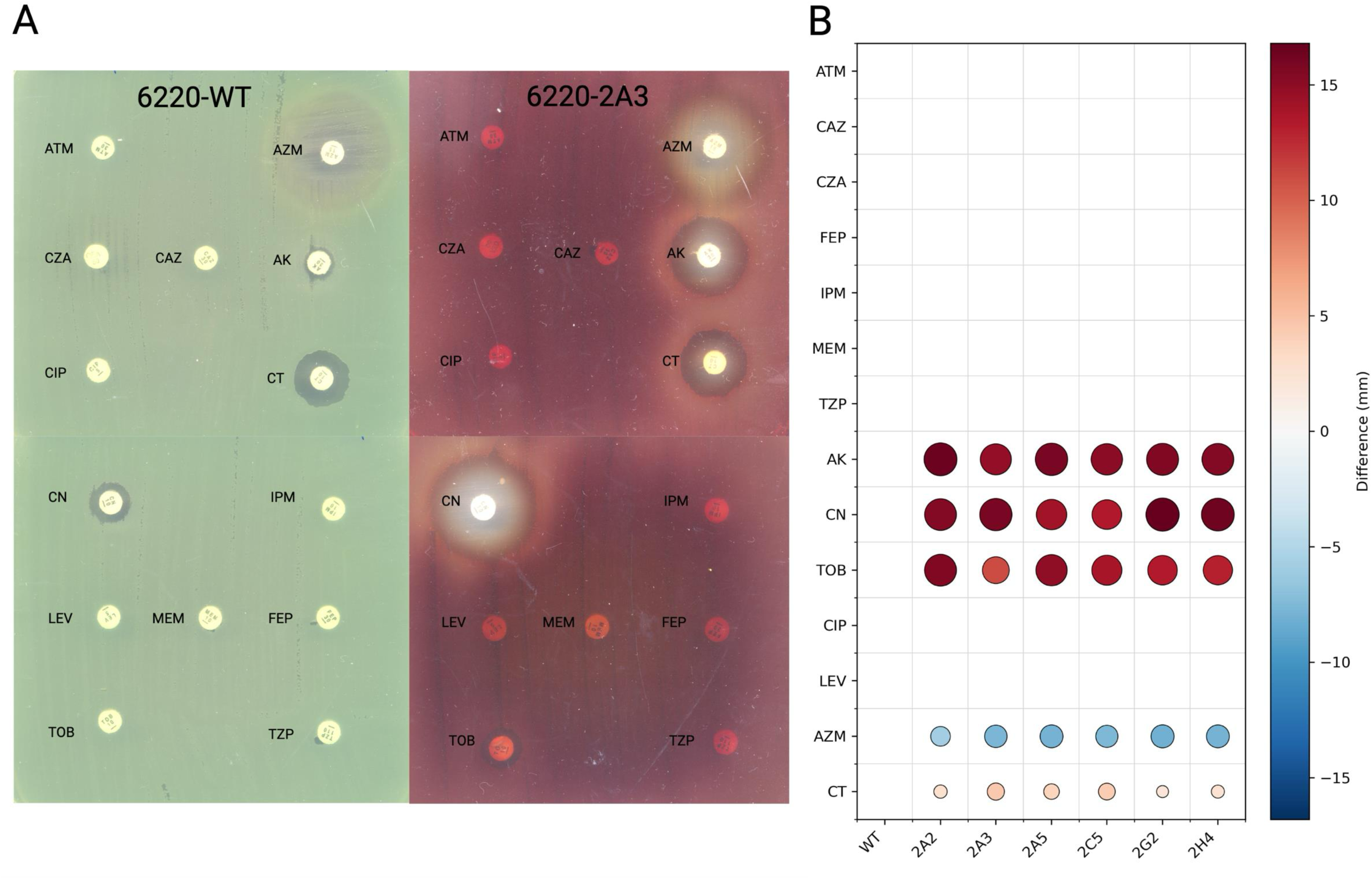
R-pyocin resistance drives a collateral sensitivity trade-off to aminoglycosides. (**A**) Kirby-Bauer disk diffusion assays showing changes in the zone of inhibition diameters for the PDR 6220-WT and a Brn mutant, 6220-2A3. Data are plotted against a panel of 14 antibiotics. (**B**) Heatmap representing the change in zone of inhibition diameter (mm) for each mutant relative to the MRSN 6220-WT ancestor and six independently generated R-pyocin-selected Brn mutants (2A2, 2A3, 2A5, 2C5, 2G2, and 2H4). Red circles indicate a positive shift, restoring sensitivity to the antibiotic, characterized by the robust sensitization of all Brn mutants to the aminoglycosides amikacin (AK), gentamicin (CN) and tobramycin (TOB). Zone diameters for colistin (CT) and azithromycin (AZM) are shown as measured values only and were not interpreted categorically, because disk diffusion does not reliably predict polymyxin susceptibility and CLSI provides no interpretive criteria for azithromycin against *P. aeruginosa*.

**Figure 5:**
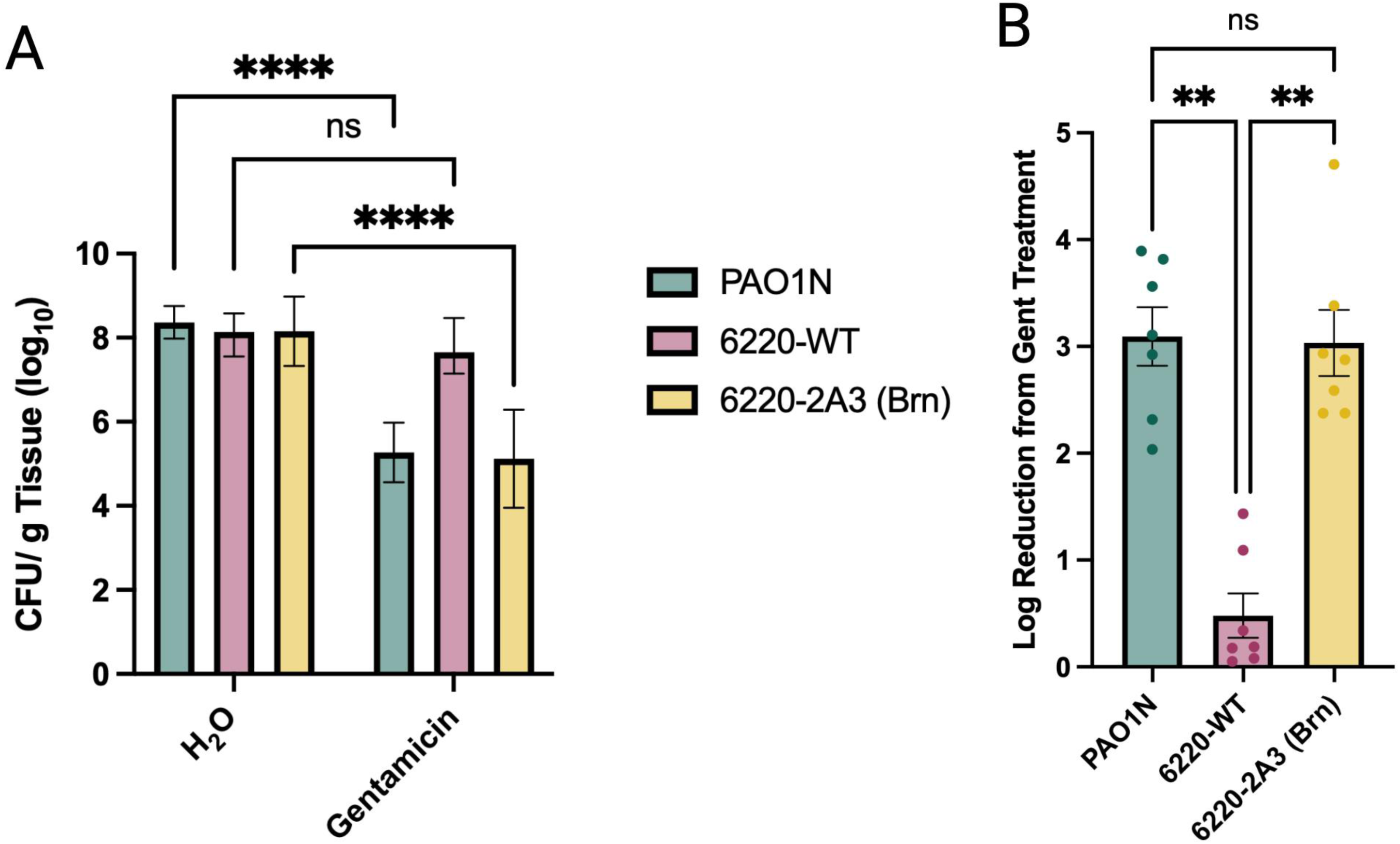
Reversion of pan-drug restores gentamicin activity against *P. aeruginosa* in a murine wound infection model. (**A**) CFU/g tissue from mouse wounds that were infected for 4 days with PAO1-WT, 6220-WT or Brn mutant 6220-2A3 and then treated ex vivo for 5 hrs with 200 µg/mL gentamicin or saline. Results reveal that the 6220-2A3 Brn mutant is sensitive to gentamicin in a complex wound environment, while the PDR ancestor 6220-WT remains refractory to the antibiotic. Statistical significance was determined using an ordinary two-way ANOVA followed by Šídák’s multiple comparisons test (n=7 per group), where mutant and reference reductions were highly significant (p < 0.0001) and the ancestral reduction was not significant (p = 0.5238). (**B**) Log reduction for gentamicin treatment for PAO1-WT, 6220-WT and 6220-2A3 Brn. Analysis confirms the selected Brn mutant achieves a significant 3-log reduction in bacterial burden comparable to the sensitive PAO1-WT control. Results represent combined data from independent replicates where n=7 for each group. Statistical significance was determined using a Kruskal-Wallis test followed by Dunn’s post-hoc multiple comparisons test. ns, not significant (p > 0.05); (**) p < 0.01; (****) p < 0.0001. Error bars represent ±SEM.

Building upon the findings of attenuated virulence and reduced aminoglycoside resistance, we evaluated the therapeutic efficacy of conventional treatment against these re-sensitized strains within a complex murine wound infection model (**Fig. 5**) (32–34). Murine wounds were infected with the PDR ancestor MRSN 6220-WT and after 4 days of infection the wound tissue was excised and treated with gentamicin. Because each variant was introduced as the inoculum, this design tests whether a previously selected mutant retains antibiotic susceptibility in a tissue environment and does not test whether R-pyocin treatment of an established infection selects the same variant *in vivo*. In this physiologically demanding tissue environment, as expected, 6220-WT remained entirely refractory to high levels of gentamicin treatment (200 µg/mL), displaying negligible reductions in bacterial load (**Fig. 5A**). In contrast, wound tissue from mice infected with the 6220-Brn mutants exhibited a robust, highly significant response to gentamicin, achieving an approximate 3-log unit reduction in bacterial burden (**Fig. 5B**). Statistical validation via a Kruskal-Wallis test (p=0.0001) with post-hoc Dunn’s multiple comparisons confirmed that this clearance was significantly greater than that observed for the treated 6220-WT ancestor (p=0.0067). Significantly, this therapeutic rescue was statistically indistinguishable from the clearance observed for the inherently sensitive PAO1 reference control (p>0.9999). Taken together, these data demonstrate that the complex physiological environment of a 4-day-old wound tissue infection does not fully shield the reverted Brn mutants from antibiotic penetration, leading to significant bactericidal rescue by gentamicin within a 5 h treatment window. Because treatment was applied to excised tissue, these data establish susceptibility at the level of infected tissue rather than therapeutic efficacy in a living host.

### A combination of R-pyocin and gentamicin suppresses growth of PDR *P. aeruginosa* and closes the pyomelanogenic escape

Because monotherapy with R-pyocins predictably drove the emergence of pyomelanogenic Brn mutants, we reasoned that combining R-pyocins with conventional aminoglycosides would close this evolutionary escape route, simultaneously suppressing the ancestral population and preventing mutant breakthrough. To test this, we performed checkerboard minimal inhibitory concentration (MIC) assays combining an R5-pyocin with gentamicin (**Fig. 6A, B**). For the laboratory reference strain PAO1-WT, the baseline MIC for gentamicin alone was 1 µg/mL. Under co-treatment, the addition of R5-pyocin at concentrations of 3.125 µg/mL and above abolished detectable growth over the 20hr assay period, and no pigmented breakthrough variants were recovered from these wells. For PDR MRSN 6220-WT, the baseline MIC for gentamicin alone was at 64 µg/mL, while the MIC for R5-pyocin alone was 6.25 µg/mL (with frequent Brn mutant breakthrough observed at higher pyocin concentrations). When the two agents were combined, a sub-lethal concentration of R5-pyocin (3.125 µg/mL) successfully potentiated gentamicin efficacy, driving the required antibiotic MIC down 8-fold to just 8 µg/mL. To quantitatively evaluate this combinatorial interaction, we calculated the Fractional Inhibitory Concentration Index (FICI) and the Dose Reduction Index (DRI). Against the PDR 6220-WT strain, the combination yielded an FICI of 0.625, indicating a strong additive-to-synergistic interaction, and a DRI value of 8, confirming a clinically meaningful 8-fold reduction in the required therapeutic dose of gentamicin. These findings demonstrate that dual-targeting strategies can exploit the evolutionary trade-offs of resistance, suppressing the growth of PDR *P. aeruginosa* and closing the pyomelanogenic escape route that R-pyocin monotherapy reliably opens, while rescuing the use of traditional, trusted antibiotics at clinically achievable concentrations. Because our endpoint measures OD_600_ rather than viability, these data define inhibition of detectable growth rather than elimination of all viable cells.

**Figure 6:**
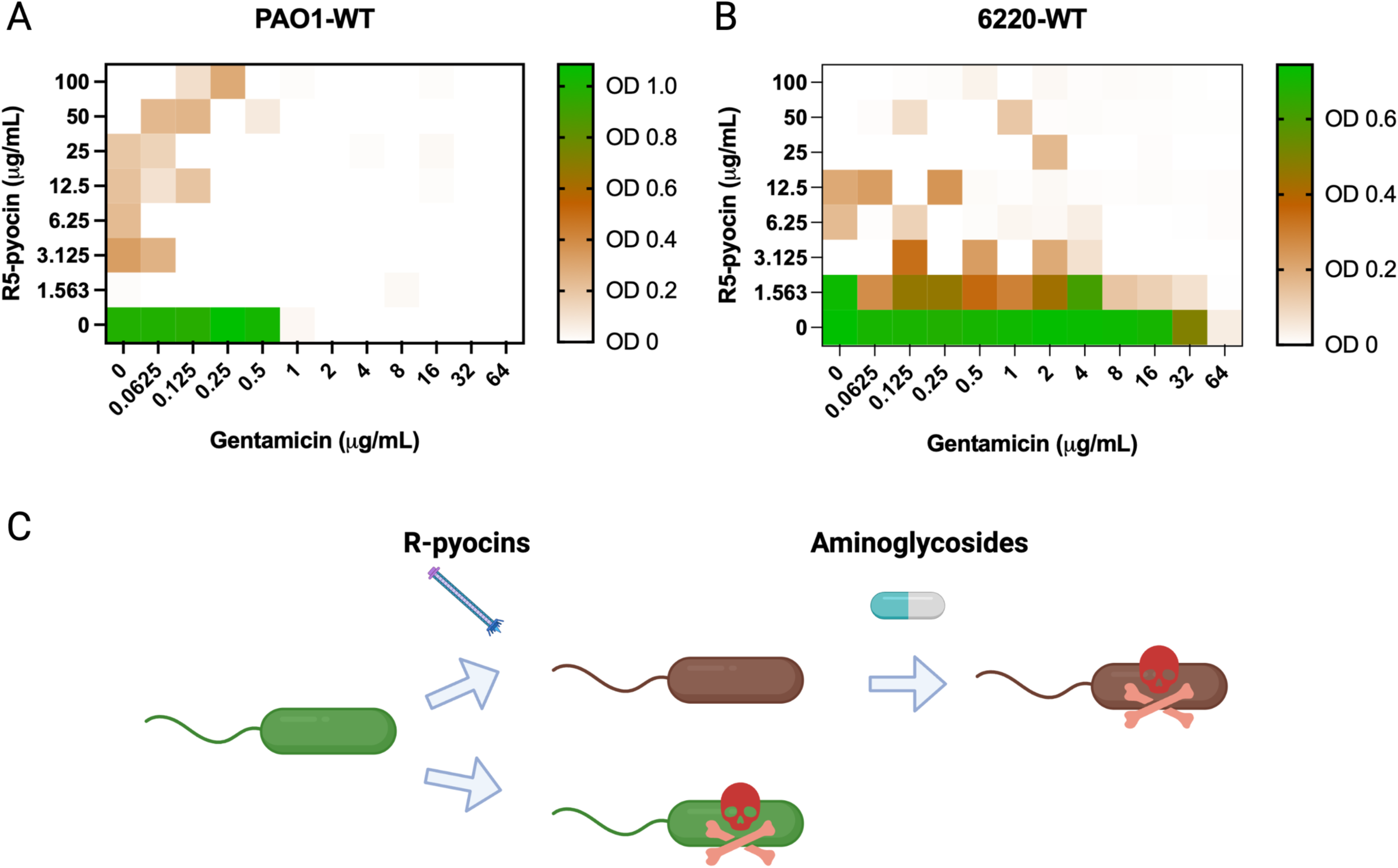
Combinatorial growth inhibition assays and mechanistic model of R-pyocin-antibiotic co-treatment. **(A)** Heatmap matrix of OD_600_ measurements of PAO1-WT exposed to a checkerboard gradient of R5-pyocins (0 to 200 µg/mL) and gentamicin (0 to 64 µg/mL). Color scale bar indicates measured bacterial density ranging from 0 (white), to 1.0 (green). **(B)** Heatmap matrix of 6220-WT under identical checkerboard treatment conditions as previously stated. **(C)** Schematic model outlining the co-treatment mechanism of R-pyocin-antibiotic combinations. Target cells (green) exposed to R-pyocins are either directly eliminated by the bactericidal activity of the tailocins or selected for as avirulent Brn mutants which are sensitized to aminoglycosides.

## Discussion

The escalating global crisis of antimicrobial resistance demands therapeutic strategies that transcend the traditional arms race of discovering new biocides. Effective next-generation interventions could also actively dismantle the pathogen’s adaptive capacity, forcing it into an evolutionary trap.

In this study, we demonstrate that R-pyocins can be deployed to engineer such an evolutionary trap in *P. aeruginosa*. By exploiting the highly conserved LPS core as a primary binding receptor (11, 13, 27, 35), R-pyocin-mediated selective pressure channels both laboratory and PDR clinical isolates toward a highly constrained genetic outcome. To survive the R-pyocin strike, the pathogen undergoes large-scale chromosomal deletions (LCDs) that trade R-pyocin resistance for collateral antibiotic sensitivity and a reduction in virulence and biofilm forming capabilities.

While the evolutionary trajectory (R-pyocin resistance coupled with a diagnostic brown pyomelanogenic phenotype driven by *hmgA* deletion) is highly conserved across diverse lineages, our complementation data reveal that the precise genetic rules governing this trade-off are remarkably strain-dependent **(Fig. 3A, B, C)**. In the clinical PDR wound isolate MRSN 6220 and laboratory strain PAK, resistance is primarily driven by the excision of *galU*, and plasmid-borne restoration of the LPS outer core fully reverts the resistant phenotype. Conversely, the failure of *galU* to complement the laboratory strain PAO1 indicates that restoration of UDP-glucose availability is not sufficient to rebuild a functional R-pyocin receptor in this background. Whether this reflects additional loci within the PAO1-specific deletion boundary, secondary changes elsewhere in the genome, or strain-level differences in LPS core structure between serotypes remains to be determined. Interestingly, the Brn phenotype and loss of *galU* has previously been found to emerge after phage exposure, with Brn mutants showing similar LCDs, but results in those studies show inconsistent, fluctuating deletion boundaries and genomic locus locations and marginal changes in antibiotic sensitivity trade-offs, often due to a reliance on working with antibiotic sensitive lab strains (37, 38). Menon et al. recovered hyperpigmented, phage-resistant PAO1 variants carrying LCDs that were more susceptible to colistin and more readily cleared by innate host defenses (39). The similarity of these findings with our own indicates that deletion-associated collateral sensitivity is a general consequence of selection by contractile particles which use the LPS core as a target, rather than a unique phenomenon to R-pyocins. Three features distinguish R-pyocin selection from phage selection. R-pyocins are non-replicative and deliver a single-hit kinetic challenge, leaving the selective pressure neither amplified by propagating phage populations nor countered by phage-encoded adaptation, which may account for the conserved deletion boundaries observed across multiple variants sourced from the same strain background and from different strain backgrounds. The collateral phenotype recovered centers on excision of the *mexXY* operon and restoration of aminoglycoside activity in a clinical isolate with a pan-drug resistant profile, rather than on polymyxin shifts in antibiotic-sensitive laboratory backgrounds. Finally, R-pyocin selection relies on purified particles rather than a replicating agent, which allows for defined dosing of the selection pressure, which is a prerequisite for the combination strategy outlined.

Crucially, the collateral damage inflicted by these LCDs extends beyond resensitizing the pathogen to frontline antibiotics. The magnitude of the genetic forfeiture forces the mutants to discard extensive virulence arsenals (hydrogen cyanide and ExoY) and major structural biofilm operons, including the *cupA1–5* fimbriae, the *bdlA* dispersal regulator, and the *fap* amyloid cluster **(Fig. 2**, **Table 2)**. This attenuation likely fundamentally alters the host-pathogen dynamic. For a patient, this evolutionary trade-off yields therapeutic advantages in which R-pyocin selection not only disarms the resistome, but the concurrent loss of structural integrity likely leaves the weakened pathogen highly vulnerable to natural host immune clearance. Without the capacity to assemble a robust, mature biofilm matrix, the pathogen is theoretically stripped of its primary structural defense against phagocytosis and neutrophil-mediated killing.

Translating these defined genetic vulnerabilities into physiological contexts will require careful model selection in the future. Our colonization data expose a critical divergence between abiotic and host-associated environments. While the LCD encompasses critical structural operons like the *cup* and *fap* clusters, and results in complete ablation of biofilm formation on plastic *in vitro* surfaces **(Fig. 3C)**, the mutants successfully achieve baseline colonization in murine wound tissue **(Fig. 5A)**. This persistence indicates that within a complex host environment, *P. aeruginosa* may be able to utilize the tissue matrix as a scaffold or rely on exopolysaccharide pathways (such as Psl/Pel or alginate) to compensate for the lost structural genes. Thus, while R-pyocin-driven selection severely compromises traditional biofilm architecture, it does not inhibit establishment of infection; highlighting the necessity of pairing R-pyocins (or phage) with traditional antibiotics to clear the pathogen.

However, the physical and biochemical complexities of the *ex vivo* tissue did not shield the reverted Brn mutants from drug penetration. A brief, 5 h gentamicin exposure was sufficient to drive significant killing of a Brn mutant when compared to the ancestor pan-drug-resistant strain (Fig. 5A, B). Whether R-pyocins reach and sustain selective concentrations within wound tissue is unknown, and diffusion through an established biofilm and necrotic matrix may generate gradients in which sub-selective exposure predominates, permitting survival without driving the deletions we describe. The net effect of these deletions on survival under oxidative stress is likewise difficult to predict. Pyomelanin overproduction, the phenotype by which we identify these mutants, confers resistance to oxidative stress and could favor the persistence of Brn variants under neutrophil attack rather than hastening their clearance, partially offsetting the attenuation conferred by loss of *hcnABC*, *exoY* and the *cupA* fimbrial genes. Working against this, the conserved deleted region also removes *katE* and *gor*, encoding catalase HPII and glutathione-disulfide reductase, so the variants that overproduce a protective pigment have simultaneously lost two enzymatic defenses against oxidative damage (Table 2). Whether Brn variants are more or less susceptible to oxidative killing than their ancestors is therefore unresolved and bears directly on whether the attenuation we observe in *Galleria* and in infected wound tissue would be reinforced or offset by the host response. Any fitness cost imposed by a 250 to 388 kbp deletion would be expected to manifest most strongly in competition with the ancestral population *in vivo*, which we have not measured. While this acute exposure model serves as a proof-of-concept at the level of infected tissue, translating this evolutionary steering into clinical practice will require extended *in vivo* animal dosing experiments to define true long-term eradication and immune clearance kinetics.

Two limitations bound the generality of this trajectory. First, our selection and screening strategy deliberately recovered pyomelanogenic variants, and the four isolates subjected to hybrid assembly were drawn from this pigmented class. Non-pigmented survivors, including those that might arise through point mutations in LPS core biosynthetic loci and would not be expected to remove mexXY, were not systematically surveyed, and we therefore cannot exclude resistance routes that avoid the collateral costs described here. Second, the frequency with which the Brn trajectory is taken relative to such alternatives remains to be quantified. The outcome we describe here is a strongly favored route under sustained R-pyocin pressure rather than an obligatory one, and the therapeutic value of the trap will depend on how reliably it is entered in a given genetic background.

Our clinical evidence also rests on multiple derivatives of a single PDR isolate, and the laboratory backgrounds, being antibiotic-sensitive at the outset, cannot extend it. We would expect the magnitude of resensitization to track the contribution of MexXY-OprM to the resistance phenotype of a given isolate, and to be correspondingly smaller in backgrounds where aminoglycoside resistance is driven by aminoglycoside-modifying enzymes or 16S rRNA methyltransferases, which lie outside the deleted region. Establishing how broadly this trade-off applies will require selection across a panel of PDR and XDR clinical isolates of differing resistance mechanism.

In summary, our findings establish R-pyocins not merely as narrow-spectrum bactericidal agents, but as potent drivers of evolutionary trajectories. The efficacy of this selective pressure is rooted in the native genomic architecture of the *P. aeruginosa* chromosome. Because loci essential for LPS core biosynthesis and R-pyocin docking, such as *galU*, are physically linked to critical multi-drug efflux systems like the *mexXY* operon, the scale of LCDs recovered here results in the co-excision of fundamental resistance determinants, and it forfeits broad-spectrum antibiotic efflux capacity. This genetic linkage establishes a strongly constrained evolutionary trade-off. By exploiting these inherent chromosomal vulnerabilities, Pyocin-Mediated Collateral Sensitivity (PMCS) provides a targeted, mechanistic strategy to dismantle the pathogen’s resistome, attenuate virulence, and restore the activity of conventional therapeutics against the pan-drug-resistant isolate examined here.

## Materials and Methods

### Bacterial strains and growth conditions

The bacterial strains utilized in this study included the pan-drug resistant *P. aeruginosa* clinical isolate MRSN 6220 (40), originally recovered from a chronic wound infection, and the laboratory strains PAO1 (Nottingham wild-type), and PAK wild-type (**Table 1**). MRSN 6220 was classified as pan-drug-resistant on the basis of non-susceptibility to all 14 agents spanning the seven antimicrobial categories used to characterize the clinical diversity panel from which it was obtained (40), a scheme adapted from Magiorakos et al. (41) in which the phosphonic acid and polymyxin categories are replaced by the β-lactam/β-lactamase inhibitor combinations ceftazidime-avibactam and ceftolozane-tazobactam.

Bacterial cultures were routinely grown in Luria-Bertani (LB) broth with constant agitation at 240 rpm, or on LB agar at 37°C. R-pyocin-resistant mutants, characterized by distinct “brown” (Brn) pigmented phenotypes, were isolated following exposure to lethal concentrations of purified R-pyocins. These variants were subsequently stabilized through multiple passages and confirmed via phenotypic retesting for pyocin resistance and pyomelanin production. For long-term storage, all strains were maintained at −80°C in LB broth supplemented with 25% (v/v) glycerol.

### R-pyocin isolation and purification

R-type pyocins were extracted as previously described (42–45), from chosen producer strains representing each subtype, R2-PAO1, and R5-1344, and two R1 variants, R1-PAK and R1-2101 (**Table 1**). R-pyocin production was induced in logarithmic-phase bacterial cultures via mitomycin C. Briefly, 500 mL of each strain was grown in LB at 37°C to an optical density (OD_600_) of ∼0.3, at which point mitomycin C (Sigma) was added to a final concentration of 3 µg/mL. Cultures were incubated with shaking for ∼3 hours until complete cell lysis occurred (as evidenced by clearing of the culture). DNase I at 1U/µL was added and incubated for an additional 30 minutes. A 1:10 ratio of chloroform was then added and incubated for another 15 minutes. Cellular debris was removed by centrifugation (4,200 ×g, 30 min, 4°C). The supernatants containing crude pyocins were collected and filtered through a 0.22 µm membrane to ensure sterility. R-pyocins in the 500 mL of crude lysates were concentrated via 4M ammonium sulfate precipitation at 1mL/min stirred on ice for 1 hour. The mixture was centrifuged (35,000 ×g, 1 hr, 4°C). The pellet was resuspended in TN50 buffer (20 mM Tris-HCl, 100 mM NaCl, pH 7.5, and centrifuge again (50,000 ×g, 1 hr, 4°C). Resuspended R-pyocins were then isolated via CsCl density gradient (gradients of 1.3, 1.4, and 1.5 g/mL CsCl in TN50 buffer) in ultracentrifuge tubes (46). Ultracentrifugation was performed at 100,000 ×g for 3 hours at 4°C in a swinging-bucket rotor. After centrifugation, a visible opalescent band corresponding to the R-pyocin particles was extracted from the gradient using a syringe. R-pyocin fractions were buffer-exchanged in 100kDa Amicon units against PBS, then filter-sterilized (0.22 µm). Purified R-pyocin preparations were stored at 4°C. Protein concentrations were determined by the Bicinchoninic acid (BCA) assay (Pierce), and purity was assessed by SDS-PAGE (which showed the expected major bands corresponding to the sheath, tube, and tail fiber proteins of the R-pyocin). To ensure absence of contaminating phage, purified preparations were exposed to UV treatment for 15 mins, then plated on lawns of a highly susceptible *P. aeruginosa* indicator strain; no individual plaques were observed, confirming that the preparations contained R-pyocins and not viable phage.

### Selection of R-pyocin resistant mutants

R-pyocin-resistant variants were generated through sustained selective pressure using a broth microdilution-based approach in sterile 96-well microtiter plates, adapted from standard minimum inhibitory concentration (MIC) protocols (47). Mid-logarithmic phase cultures of *P. aeruginosa* 6220-WT, PAO1, and PAK were adjusted to a final inoculum of approximately ∼5×10⁵ CFU/mL in Cation adjusted Mueller Hinton broth and challenged with a range of concentrations (1:2 dilution) of purified R1, R2, or R5 pyocins. The plates were incubated statically at 37°C for 24 to 48 hrs, during which time the wells were monitored for persistent growth and significant phenotypic shifts. Surviving populations from wells containing R-pyocin concentrations at or above the inhibitory threshold were screened for the emergence of distinct pigmented variants, specifically identifying pyomelanogenic Brn mutants. Individual colonies were subsequently isolated by streaking onto Pseudomonas Isolation agar (PIA) plates to ensure the stability of the resistant phenotype, which was further validated through high-titer R-pyocin challenge and standardized spot assays (26, 27, 48).

### Construction of complementation strains

For genetic complementation, wild-type copies of the *galU* (49) and *hmgA* genes were introduced into the R-pyocin-resistant mutant backgrounds on a pUCP18 plasmid with a tetracycline resistance marker. The genes were expressed from plasmid vectors to evaluate the restoration of R-pyocin susceptibility via *galU* complementation, and the reversal of the pyomelanogenic pigmentation via *hmgA* complementation. Susceptibility to R-pyocin subtypes R1, R2, and R5 was subsequently re-evaluated in these complemented strains to confirm the restoration of R-pyocin susceptibility.

### Antibiotic sensitivity testing

Collateral sensitivity in R-pyocin-resistant mutants was determined using the Kirby-Bauer disk diffusion method according to standard clinical protocols (29, 30). The PDR MRSN 6220-WT ancestor and its Brn mutants were screened against a broad panel of clinically relevant antibiotics, including β-lactams (aztreonam, ceftazidime, ceftazidime-avibactam, cefepime, imipenem, meropenem, and piperacillin-tazobactam) and aminoglycosides (amikacin, gentamicin, and tobramycin). Following an 18 h incubation at 37°C, the diameters of the zones of inhibition were measured in millimeters. Zone diameters were interpreted according to the breakpoints for *P. aeruginosa* in CLSI M100, 36th edition (31), and both the measured diameter and the resulting categorical interpretation are reported for each agent in **Table S6**. Several agents in the panel cannot be assigned a susceptibility category under current criteria. CLSI does not provide interpretive breakpoints for gentamicin or azithromycin against *P. aeruginosa* and restricts amikacin interpretation to urinary isolates, and disk diffusion does not reliably predict polymyxin susceptibility. Zone diameters for these four agents are therefore reported as measured values only and are not interpreted categorically.

### Biofilm formation and *in vivo* virulence assays

Biofilm formation was quantified using the crystal violet staining method, (50) in 96-well polystyrene plates to assess the impact of *galU* deficiency and subsequent restoration via complementation. The pathogenic potential of wild-type and mutant strains was further evaluated in a *Galleria mellonella* infection model, as previously described (51). Larvae were injected with standardized bacterial suspensions, and survival and melanization were monitored over a 24-hour period post-injection. PBS-injected and no-treatment groups were included as negative controls to ensure larval viability.

### Comparative genomic analysis

Genomic architecture was resolved using a hybrid assembly approach combining long-read and short-read sequencing data (52). High-molecular-weight genomic DNA was extracted using the QIAamp DNA Blood Mini Kit (Qiagen, Germantown, MD), with concentration and purity confirmed via Qubit dsDNA Broad Range Assay (ThermoFisher Scientific) and Nanodrop 2000 spectrophotometry. For long-read sequencing, libraries were prepared using the Oxford Nanopore Technologies (ONT) Rapid Barcoding Sequencing Kit (SQK-RBK004) and sequenced on an R9.4.1 flow cell. Long-read sequences were basecalled using Guppy v.3.0.7, followed by demultiplexing and adapter trimming with Porechop v.0.2.3. Long reads were further quality-filtered and refined using Filtlong v.0.2.0, utilizing the corresponding processed short reads as a reference for quality weighting. Filtering parameters were set to a minimum length threshold of 1,000 bp, a target output of 500 Mbp, and the top 90% of quality-weighted bases, with active read trimming and splitting (--trim --split 500). BBDuk was employed for the removal of phiX and adapter sequences using reference-specific FASTA files. Subsequent quality trimming and filtering were performed with fastp; briefly, the first 15 bases were trimmed from the 5’ end of each read, poly-X tails were removed, reads shorter than 50 bp were discarded, and reads with an overall average quality score below Q30 were removed. *De novo* short-read assemblies were generated using SPAdes v.3.15.3 with the --isolates parameters. Final hybrid genome assemblies were generated using Unicycler v.0.4.8 with default parameters. Assemblies were annotated with the BAKTA Pipeline. Chromosomal deletions were detected via whole-genome alignments utilizing MUMmer/nucmer, and genomic coordinates of the structural variations were extracted using the show-coords -rcl command. The resulting architectural profiles and large-scale deletions were visualized with BLASTN, Easyfig, Pycirculize, and Matplotlib.

### Murine wound infection model

To evaluate the clinical efficacy of conventional antibiotic treatment against re-sensitized strains, a murine wound infection model was used. Briefly, 21 mice were anesthetized, the dorsum was shaved, depilated, and administered a circular, full-thickness dermal excision 1.5 cm in diameter. Excised wounds were covered with a semipermeable dressing (Opsite dressing; Smith and Nephew) and infected with a standardized inoculum of 10^4^ CFU/mL of either PAO1-WT, the PDR ancestor 6220-WT, or the R-pyocin-resistant Brn mutant 6220-2A3. After 4 days of infection, mice were euthanized and the wound beds were excised and divided, then submerged for 5 hrs in either 1 mL of sterile H_2_O or 200 µg/mL of Gentamicin. Following treatment, period, treatment solution was discarded, replaced with 1X PBS and homogenized in a bead mill (FastPrep-24™ MP Biomedicals) at 5 m/s for 120 seconds. Homogenate was pelleted (12,000 x G for 5 min) and resuspended in 1X PBS three times to remove any remaining antibacterial agent. Washed homogenate was serially diluted and plated on PIA to quantify remaining bacterial burden. Statistical significance was determined using a Kruskal-Wallis test followed by Dunn’s multiple comparisons test to evaluate differences in log reduction between groups.

### Combinatorial checkerboard broth microdilution assay

To evaluate the combinatorial efficacy of R-pyocins and conventional aminoglycosides against both PAO1-WT and a MRSN 6220-WT, checkerboard broth microdilution assays were performed in 96-well flat-bottom plates (53). Bacterial inoculum was prepared from overnight cultures and adjusted to an OD_600_ ∼0.8, then diluted 1:200 in cation-adjusted Mueller_Hinton broth (CAMHB) to achieve a starting concentration of ∼5×10⁵ CFU/mL. Two-fold serial dilutions of gentamicin sulfate were prepared in 0.85% saline across columns, and a two-fold serial dilution of R5-pyocin were prepared in TN50 buffer across rows using separate master plates. Each experimental well received 50 mL of the respective R-pyocin dilution and 50 mL of the gentamicin dilution, followed by 100 mL of the bacterial inoculum. Parallel control plates were prepared, including uninoculated background control wells containing CAMHB, R-pyocins, and saline to account for R-pyocin opacity, along with standard broth sterility controls. Plates were incubated statically at 37°C for 15-20 hours. Following incubation, endpoint was determined by visual inspection and quantified be OD_600_ measurement. The minimum inhibitory concentration was defined as the lowest concentration that completely inhibited visible bacterial growth. Growth in each well was scored by visual inspection and OD_600_measurement. This endpoint reports inhibition of detectable growth and does not distinguish bacteriostatic from bactericidal activity. To characterize the interaction profile, the Fractional Inhibitory Concentration (FIC) index was calculated for wells demonstrating complete growth inhibition using the FICI formula: 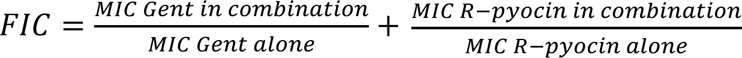. Additionally, the Dose Reduction Index was calculated using the Chou-Talalay formula: 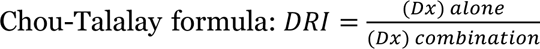

## Acknowledgements

For funding, we thank the Georgia Institute of Technology and the National Institutes of Health for grants (NIAID R56AI184449; R01AI153116) to S.P.D, NIH (R21AI178702) to J.B.G, the Cystic Fibrosis Foundation (GOLDBE23G0; DIGGLE26G0) to J.B.G and S.P.D., and F31 AI191664-02 to I.E.

## Author contributions

All authors contributed to research design; I.E., G.W., P.S., and D.C. performed research and analyzed data; all authors contributed to writing the paper. All figures created in BioRender. Estrada, I. (2026)

## References

1. Qin S, Xiao W, Zhou C, Pu Q, Deng X, Lan L, Liang H, Song X, Wu M. 2022. Pseudomonas aeruginosa: pathogenesis, virulence factors, antibiotic resistance, interaction with host, technology advances and emerging therapeutics. 1. Signal Transduct Target Ther 7:1–27.

2. Letizia M, Diggle SP, Whiteley M. 2025. Pseudomonas aeruginosa: ecology, evolution, pathogenesis and antimicrobial susceptibility. Nat Rev Microbiol 23:701–717.

3. Hernandez-Beltran JCR, Rodríguez-Beltrán J, Aguilar-Luviano OB, Velez-Santiago J, Mondragón-Palomino O, MacLean RC, Fuentes-Hernández A, San Millán A, Peña-Miller R. 2024. Plasmid-mediated phenotypic noise leads to transient antibiotic resistance in bacteria. Nat Commun 15:2610.

4. Pang Z, Raudonis R, Glick BR, Lin T-J, Cheng Z. 2019. Antibiotic resistance in *Pseudomonas aeruginosa*: mechanisms and alternative therapeutic strategies. Biotechnol Adv 37:177–192.

5. López-Causapé C, Cabot G, del Barrio-Tofiño E, Oliver A. 2018. The Versatile Mutational Resistome of Pseudomonas aeruginosa. Front Microbiol 9:685.

6. Khuntayaporn P, Yamprayoonswat W, Yasawong M, Chomnawang MT. 2019. Dissemination of Carbapenem-Resistance among Multidrug Resistant Pseudomonas aeruginosa carrying Metallo-Beta-Lactamase Genes, including the Novel blaIMP-65 Gene in Thailand. Infect Chemother 51:107–118.

7. Morita Y, Tomida J, Kawamura Y. 2012. MexXY multidrug efflux system of Pseudomonas aeruginosa. Front Microbiol 3:408.

8. Fujiwara M, Yamasaki S, Morita Y, Nishino K. 2022. Evaluation of efflux pump inhibitors of MexAB- or MexXY-OprM in *Pseudomonas aeruginosa* using nucleic acid dyes. J Infect Chemother 28:595–601.

9. Mei M, Estrada I, Diggle SP, Goldberg JB. 2025. R-pyocins as targeted antimicrobials against Pseudomonas aeruginosa. Npj Antimicrob Resist 3:17.

10. Estrada I, Smith P, Mei M, Goldberg JB, Diggle SP. 2025. Microbial Primer: The R-pyocins of Pseudomonas aeruginosa. Microbiology 171:001640.

11. Köhler T, Donner V, van Delden C. 2010. Lipopolysaccharide as Shield and Receptor for R-Pyocin-Mediated Killing in Pseudomonas aeruginosa. J Bacteriol 192:1921–1928.

12. Nakayama K, Takashima K, Ishihara H, Shinomiya T, Kageyama M, Kanaya S, Ohnishi M, Murata T, Mori H, Hayashi T. 2000. The R-type pyocin of *Pseudomonas aeruginosa* is related to P2 phage, and the F-type is related to lambda phage. Mol Microbiol 38:213–231.

13. Michel-Briand Y, Baysse C. 2002. The pyocins of Pseudomonas aeruginosa. Biochimie 84:499–510.

14. Lee FKN, Dudas KC, Hanson JA, Nelson MB, LoVerde PT, Apicella MA. 1999. The R-Type Pyocin of Pseudomonas aeruginosa C Is a Bacteriophage Tail-Like Particle That Contains Single-Stranded DNA. Infect Immun 67:717–725.

15. Ge P, Scholl D, Prokhorov NS, Shneider MM, Browning C, Buth SA, Plattner M, Chakraborty U, Ding K, Leiman PG, Miller JF, Zhou ZH. 2020. Action of a minimal contractile bactericidal nanomachine. 7805. Nature 580:658–662.

16. Patz S, Becker Y, Richert-Pöggeler KR, Berger B, Ruppel S, Huson DH, Becker M. 2019. Phage tail-like particles are versatile bacterial nanomachines – A mini-review. J Adv Res 19:75–84.

17. Ge P, Scholl D, Leiman PG, Yu X, Miller JF, Zhou ZH. 2015. Atomic structures of a bactericidal contractile nanotube in its pre- and postcontraction states. Nat Struct Mol Biol 22:377–382.

18. Kageyama M. 1964. Studies of a Pyocin: I. Physical and Chemical Properties*. J Biochem (Tokyo) 55:49–53.

19. Kageyama M, Ikeda K, Egami F. 1964. Studies of a Pyocin: III. Biological Properties of the Pyocin*. J Biochem (Tokyo) 55:59–64.

20. Kageyama M, Egami F. 1962. On the purification and some properties of a pyocin, a bacteriocin produced by Pseudomonas aeruginosa. Life Sci 1:471–476.

21. Bae H-W, Choi S-Y, Ki H-J, Cho Y-H. 2025. Pseudomonas aeruginosa as a model bacterium in antiphage defense research. FEMS Microbiol Rev 49:fuaf014.

22. Costa AR, van den Berg DF, Esser JQ, Muralidharan A, van den Bossche H, Bonilla BE, van der Steen BA, Haagsma AC, Fluit AC, Nobrega FL, Haas P-J, Brouns SJJ. 2024. Accumulation of defense systems in phage-resistant strains of Pseudomonas aeruginosa. Sci Adv 10:eadj0341.

23. Burke KA, Urick CD, Mzhavia N, Nikolich MP, Filippov AA. 2024. Correlation of Pseudomonas aeruginosa Phage Resistance with the Numbers and Types of Antiphage Systems. Int J Mol Sci 25:1424.

24. Shen M, Zhang H, Shen W, Zou Z, Lu S, Li G, He X, Agnello M, Shi W, Hu F, Le S. 2018. Pseudomonas aeruginosa MutL promotes large chromosomal deletions through non-homologous end joining to prevent bacteriophage predation. Nucleic Acids Res 46:4505–4514.

25. Oliver A, Baquero F, Blázquez J. 2002. The mismatch repair system (mutS, mutL and uvrD genes) in Pseudomonas aeruginosa: molecular characterization of naturally occurring mutants. Mol Microbiol 43:1641–1650.

26. Mei M, Thomas J, Diggle SP. 2021. Heterogenous Susceptibility to R-Pyocins in Populations of Pseudomonas aeruginosa Sourced from Cystic Fibrosis Lungs. mBio 12:e00458–21.

27. Mei M, Pheng P, Kurzeja-Edwards D, Diggle SP. 2023. High prevalence of lipopolysaccharide mutants and R2-pyocin susceptible variants in Pseudomonas aeruginosa populations sourced from cystic fibrosis lung infections. Microbiol Spectr 11:e0177323.

28. Secor PR, Sweere JM, Michaels LA, Malkovskiy AV, Lazzareschi D, Katznelson E, Rajadas J, Birnbaum ME, Arrigoni A, Braun KR, Evanko SP, Stevens DA, Kaminsky W, Singh PK, Parks WC, Bollyky PL. 2015. Filamentous Bacteriophage Promote Biofilm Assembly and Function. Cell Host Microbe 18:549–559.

29. Schiller H, Young C, Schulze S, Tripepi M, Pohlschroder M. A Twist to the Kirby-Bauer Disk Diffusion Susceptibility Test: an Accessible Laboratory Experiment Comparing Haloferax volcanii and Escherichia coli Antibiotic Susceptibility to Highlight the Unique Cell Biology of Archaea. J Microbiol Biol Educ 23:e00234–21.

30. Biemer JJ. 1973. Antimicrobial Susceptibility Testing by the Kirby-Bauer Disc Diffusion Method. Ann Clin Lab Sci 3:135–140.

31. Clinical and Laboratory Standards Institute. 2026. M100 | Performance Standards for Antimicrobial Susceptibility Testing, 36th ed. Clinical and Laboratory Standards Institute, Wayne, PA. https://clsi.org/shop/standards/m100/. Retrieved 23 June 2026.

32. Fleming D, Chahin L, Rumbaugh K. 2017. Glycoside Hydrolases Degrade Polymicrobial Bacterial Biofilms in Wounds. Antimicrob Agents Chemother 61:10.1128/aac.01998-16.

33. Redman WK, Welch GS, Rumbaugh KP. 2020. Differential Efficacy of Glycoside Hydrolases to Disperse Biofilms. Front Cell Infect Microbiol 10.

34. Vanderwoude J, Fleming D, Azimi S, Trivedi U, Rumbaugh KP, Diggle SP. 2020. The evolution of virulence in Pseudomonas aeruginosa during chronic wound infection. Proc R Soc B Biol Sci 287:20202272.

35. Scholl D. 2017. Phage Tail–Like Bacteriocins. Annu Rev Virol 4:453–467.

36. Nowicki EM, O’Brien JP, Brodbelt JS, Trent MS. 2015. Extracellular zinc induces phosphoethanolamine addition to Pseudomonas aeruginosa lipid A via the ColRS two-component system. Mol Microbiol 97:166–178.

37. Nakamura K, Fujiki J, Nakamura T, Furusawa T, Gondaira S, Usui M, Higuchi H, Tamura Y, Iwano H. 2021. Fluctuating Bacteriophage-induced galU Deficiency Region is Involved in Trade-off Effects on the Phage and Fluoroquinolone Sensitivity in Pseudomonas aeruginosa. Virus Res 306:198596.

38. Fujiki J, Nakamura K, Ishiguro Y, Iwano H. 2024. Using phage to drive selections toward restoring antibiotic sensitivity in Pseudomonas aeruginosa via chromosomal deletions. Front Microbiol 15.

39. Menon ND, Penziner S, Montaño ET, Zurich R, Pride DT, Nair BG, Kumar GB, Nizet V. 2022. Increased Innate Immune Susceptibility in Hyperpigmented Bacteriophage-Resistant Mutants of Pseudomonas aeruginosa. Antimicrob Agents Chemother 66:e0023922.

40. Lebreton F, Snesrud E, Hall L, Mills E, Galac M, Stam J, Ong A, Maybank R, Kwak YI, Johnson S, Julius M, Ly M, Swierczewski B, Waterman PE, Hinkle M, Jones A, Lesho E, Bennett JW, McGann P. 2021. A panel of diverse Pseudomonas aeruginosa clinical isolates for research and development. JAC-Antimicrob Resist 3:dlab179.

41. Magiorakos A-P, Srinivasan A, Carey RB, Carmeli Y, Falagas ME, Giske CG, Harbarth S, Hindler JF, Kahlmeter G, Olsson-Liljequist B, Paterson DL, Rice LB, Stelling J, Struelens MJ, Vatopoulos A, Weber JT, Monnet DL. 2012. Multidrug-resistant, extensively drug-resistant and pandrug-resistant bacteria: an international expert proposal for interim standard definitions for acquired resistance. Clin Microbiol Infect Off Publ Eur Soc Clin Microbiol Infect Dis 18:268–281.

42. Scholl D, Cooley M, Williams SR, Gebhart D, Martin D, Bates A, Mandrell R. 2009. An Engineered R-Type Pyocin Is a Highly Specific and Sensitive Bactericidal Agent for the Food-Borne Pathogen Escherichia coli O157:H7. Antimicrob Agents Chemother 53:3074–3080.

43. Ibarguren-Quiles C, Blasco L, López-Causape C, Bleriot I, Fernández-García L, Arman L, Barrio-Pujante A, Ortiz-Cartagena C, Aracil B, Menéndez-Rodriguez O, Mariñas-Pardo L, Cantón R, Oliver A, Tomás M. 2026. Identification and functional insights into new phage tail-like bacteriocins targeting Pseudomonas aeruginosa as new antimicrobials. Microbiol Spectr 0:e02894–25.

44. Alqahtani A, Kopel J, Hamood A. 2022. The In Vivo and In Vitro Assessment of Pyocins in Treating Pseudomonas aeruginosa Infections. Antibiot Basel Switz 11:1366.

45. Scholl D, Martin DW. 2008. Antibacterial Efficacy of R-Type Pyocins towards Pseudomonas aeruginosa in a Murine Peritonitis Model. Antimicrob Agents Chemother 52:1647–1652.

46. Lab JK. 2016. CsCl Step Gradient to Purify Phage.

47. Kadeřábková N, Mahmood AJS, Mavridou DAI. 2024. Antibiotic susceptibility testing using minimum inhibitory concentration (MIC) assays. Npj Antimicrob Resist 2:37.

48. Oluyombo O, Penfold CN, Diggle SP. 2019. Competition in Biofilms between Cystic Fibrosis Isolates of Pseudomonas aeruginosa Is Shaped by R-Pyocins. mBio 10:10.1128/mbio.01828-18.

49. Dean CR, Goldberg JB. 2002. Pseudomonas aeruginosa galU is required for a complete lipopolysaccharide core and repairs a secondary mutation in a PA103 (serogroup O11) wbpM mutant. FEMS Microbiol Lett 210:277–283.

50. Feoktistova M, Geserick P, Leverkus M. 2016. Crystal Violet Assay for Determining Viability of Cultured Cells. Cold Spring Harb Protoc 2016:pdb.prot087379.

51. Berryhill BA, Gil-Gil T, Witzany C, Goldberg DA, Vega NM, Regoes RR, Levin BR. 2025. The role of innate immunity, antibiotics, and bacteriophages in the course of bacterial infections and their treatment. Proc Natl Acad Sci 122:e2507914122.

52. Kent AG, Spicer LM, Campbell D, Breaker E, McAllister GA, Ewing TO, Longo C, Balbuena R, Burroughs M, Burgin A, Padilla J, Johnson JK, Halpin AL, McKay SL, Rasheed JK, Elkins CA, Karlsson M, Lutgring JD, Gargis AS. Sentinel Surveillance reveals phylogenetic diversity and detection of linear plasmids harboring vanA and optrA among enterococci collected in the United States. Antimicrob Agents Chemother 68:e00591–24.

53. Bellio P, Fagnani L, Nazzicone L, Celenza G. 2021. New and simplified method for drug combination studies by checkerboard assay. MethodsX 8:101543.

54. Fractional Inhibitory Concentration Index - an overview | ScienceDirect Topics. https://www.sciencedirect.com/topics/immunology-and-microbiology/fractional-inhibitory-concentration-index. Retrieved 28 May 2026.

55. Chou T-C. 2010. Drug Combination Studies and Their Synergy Quantification Using the Chou-Talalay Method. Cancer Res 70:440–446.

