## Supplementary material for "R-pyocin-mediated selection restores aminoglycoside sensitivity in a pan-drug resistant *Pseudomonas aeruginosa* isolate": Supp Table S6

**Table S6.** Zone diameters (mm) and CLSI categorical interpretations for MRSN 6220-WT and six independently selected R-pyocin-resistant Brn mutants.

Antimicrobial categories follow the seven-category scheme used to characterize the clinical diversity panel from which MRSN 6220 was obtained, adapted from Magiorakos et al. by replacing the phosphonic acid and polymyxin categories with the β-lactam/β-lactamase inhibitor combinations. Values are given as zone diameter in mm followed by the susceptibility category in parentheses. S, susceptible; I, intermediate; R, resistant; NI, not interpreted.

| **Antimicrobial category** | **Antimicrobial agent** | **Abbrev.** | **6220-WT** | **6220-2A2** | **6220-2A3** | **6220-2A5** | **6220-2C5** | **6220-2G2** | **6220-2H4** |
| --- | --- | --- | --- | --- | --- | --- | --- | --- | --- |
| Aminoglycosides | Amikacin | AK | 7.3 (NI) | 23.8 (NI) | 22 (NI) | 23.3 (NI) | 22.3 (NI) | 22.9 (NI) | 22.7 (NI) |
|  | Gentamicin | CN | 10.4 (NI) | 25.8 (NI) | 26.3 (NI) | 24.5 (NI) | 23.8 (NI) | 27.2 (NI) | 26.8 (NI) |
|  | Tobramycin | TOB | 0 (R) | 15.5 (I) | 11 (R) | 14.9 (I) | 13.8 (I) | 13.3 (I) | 13.1 (I) |
| Antipseudomonal carbapenems | Imipenem | IPM | 0 (R) | 0 (R) | 0 (R) | 0 (R) | 0 (R) | 0 (R) | 0 (R) |
|  | Meropenem | MEM | 0 (R) | 0 (R) | 0 (R) | 0 (R) | 0 (R) | 0 (R) | 0 (R) |
| Antipseudomonal cephalosporins | Ceftazidime | CAZ | 0 (R) | 0 (R) | 0 (R) | 0 (R) | 0 (R) | 0 (R) | 0 (R) |
|  | Cefepime | FEP | 0 (R) | 0 (R) | 0 (R) | 0 (R) | 0 (R) | 0 (R) | 0 (R) |
| Antipseudomonal fluoroquinolones | Ciprofloxacin | CIP | 0 (R) | 0 (R) | 0 (R) | 0 (R) | 0 (R) | 0 (R) | 0 (R) |
|  | Levofloxacin | LEV | 0 (R) | 0 (R) | 0 (R) | 0 (R) | 0 (R) | 0 (R) | 0 (R) |
| Antipseudomonal penicillins + β-lactamase inhibitors | Piperacillin-tazobactam | TZP | 0 (R) | 0 (R) | 0 (R) | 0 (R) | 0 (R) | 0 (R) | 0 (R) |
| Monobactams | Aztreonam | ATM | 0 (R) | 0 (R) | 0 (R) | 0 (R) | 0 (R) | 0 (R) | 0 (R) |
| β-lactam/β-lactamase inhibitor combinations | Ceftazidime-avibactam | CZA | 0 (R) | 0 (R) | 0 (R) | 0 (R) | 0 (R) | 0 (R) | 0 (R) |
| Not represented in the categorization scheme | Azithromycin | AZM | 33.8 (NI) | 27.9 (NI) | 26.1 (NI) | 25.8 (NI) | 26.3 (NI) | 25.7 (NI) | 25.8 (NI) |
|  | Colistin | CT | 14.6 (NI) | 17.3 (NI) | 19.1 (NI) | 18.3 (NI) | 18.9 (NI) | 16.8 (NI) | 17.2 (NI) |

**a** Zone diameters are reported as the mean of [— N] independent biological replicates. A value of 0 indicates no measurable zone of inhibition.

**b** Amikacin. CLSI restricts interpretation of amikacin against P. aeruginosa to urinary isolates. MRSN 6220 is a wound isolate, so amikacin zone diameters are reported as measured values and are not interpreted categorically.

**c** Gentamicin. CLSI no longer provides interpretive criteria for gentamicin against P. aeruginosa. Zone diameters are reported as measured values and are not interpreted categorically.

**d** Tobramycin. Interpreted using the CLSI M100 criteria for P. aeruginosa (S ≥ 19 mm, I 13-18 mm, R ≤ 12 mm). Five of six Brn mutants moved from the resistant into the intermediate category; none reached susceptible.

**e** Azithromycin. CLSI provides no interpretive criteria for azithromycin against P. aeruginosa. Zone diameters are reported as measured values and are not interpreted categorically.

**f** Colistin. Disk diffusion does not reliably predict polymyxin susceptibility and is not a valid method for colistin. Zone diameters are reported as measured values and are not interpreted categorically. Because the polymyxin category is not represented in the categorization scheme applied here, the pan-drug-resistant designation of MRSN 6220 carries no claim regarding colistin susceptibility.

**g** Agents marked R at 0 mm are resistant by any current CLSI criterion, as no measurable inhibition was observed.

**h** Ticarcillin-clavulanic acid and ceftolozane-tazobactam, the second agents in the antipseudomonal penicillin and β-lactam/β-lactamase inhibitor categories respectively, were not included in this panel. All seven antimicrobial categories are nonetheless represented.

**Summary of the resistance profile supporting the PDR designation**

MRSN 6220-WT was non-susceptible to every agent tested across all seven antimicrobial categories. Of the 14 agents in this panel, 12 fall within the categorization scheme and 2 (azithromycin and colistin) lie outside it. Ticarcillin-clavulanic acid and ceftolozane-tazobactam were not tested, but each of their categories is represented by another agent.

*Verify the tobramycin and amikacin criteria against CLSI M100, 36th edition, Table 2B-2 before submission. The values applied here are drawn from the breakpoint revisions introduced in M100 ed. 33 and reported in the FDA recognition decision for the aminoglycosides.*
